# PopB-PcrV Interactions are Essential for Pore Formation in the *Pseudomonas aeruginosa* Type III Secretion System Translocon

**DOI:** 10.1101/2022.01.21.477233

**Authors:** Emma Caitlin Kundracik, Josephine Trichka, José Enrique Díaz Aponte, Arne Rietsch

**Affiliations:** Dept. of Molecular Biology and Microbiology, Case Western Reserve University, Cleveland, OH, U.S.A.; Dept. of Pathology, Case Western Reserve University, Cleveland, OH, U.S.A.; Dept. of Physiology and Biophysics, Case Western Reserve University, Cleveland, OH, U.S.A.

**Keywords:** T3SS, translocation, pore formation

## Abstract

The type III secretion system (T3SS) is a syringe-like virulence factor which delivers bacterial proteins directly into the cytoplasm of host cells. An essential component of the system is the translocon, which creates a pore in the host cell membrane through which proteins are injected. In *Pseudomonas aeruginosa*, the translocation pore is formed by proteins PopB and PopD and attaches to the T3SS needle via the needle tip protein PcrV. The pore is multimeric, but the exact stoichiometry and structure of the pore are unknown. We took a genetic approach to map contact points within the system by taking advantage of the fact that the translocator proteins of *Pseudomonas aeruginosa* and the related *Aeromonas hydrophila* T3SS are incompatible and cannot be freely exchanged. We created chimeric versions of *P. aeruginosa* PopB and *A. hydrophila* AopB to intentionally disrupt and restore protein-protein interactions. We identified a chimeric B-translocator that specifically breaks an interaction with the needle tip protein and interferes with the formation of the translocation pore. Breaking the interaction did not disrupt membrane insertion, arguing that the needle tip protein chaperones formation of the translocation pore.

## Introduction

The type III secretion system (T3SS) is a prime target for anti-virulence drug development against multidrug resistant *Pseudomonas aeruginosa* (1–3). This syringe-like virulence factor is used by *P. aeruginosa* to inject host cells with up to four effector proteins: ExoS, ExoT, ExoU, and ExoY. These effectors are targeted to neutrophils and macrophages to hinder immune clearance or into epithelial cells to enhance dissemination of the bacteria (4). The portion of the T3SS which creates a pore in the host cell membrane, the translocon, is an attractive therapeutic target because it is located in the extracellular space and is immunogenic (5–7). However, the translocon is poorly understood due to the many challenges presented by its unique assembly process in the host cell membrane (8, 9).

The structure of the translocon, its manner of assembly, and how it regulates effector secretion all remain open questions. Complex processes, such as the needle-tip mediated insertion of translocator proteins into the host cell membrane or triggering of effector secretion, cannot be recapitulated in vitro. The translocon readily dissociates, so the intact structure has yet to be purified. The *Salmonella* translocon was visualized in the context of an infected cell by cryo-electron tomography, but the structure lacked molecular resolution (10). A further complication, in the case of *P. aeruginosa*, is that the copy number of the assembled system is low—just 1-3 per bacterium—hampering direc*t in situ* imaging studies (11, 12). For these reasons, the structure of the translocon remains elusive despite keen interest by the T3SS research community.

Formation of the translocation pore has been studied in vitro using purified proteins (13–16). In the case of *P. aeruginosa*, two transmembrane proteins, PopB and PopD, oligomerize to form a pore in the host cell membrane. PopB and PopD can also form pores in lipid bilayers (17–19) and under the right conditions, these pores require both translocator proteins, just like translocation requires PopB and PopD in the context of host cells. More recently, the stoichiometry of translocator proteins in in vitro-formed pores was estimated using photobleaching, suggesting that these pores consist of 8 copies of PopB and 8 copies of PopD (20). Genetic studies have also shed light on the composition and function of the translocon. The needle tip protein is required for translocator insertion in every T3SS examined to date (21–26). Indeed, the *P. aeruginosa* needle tip protein PcrV connects the translocation pore to the T3SS needle and is needed for insertion of PopB and PopD into the host cell membrane. Recent data suggests that insertion of PopD is further facilitated by PopB (17). The needle tip protein plays a role in sensing host cell contact, so that effector proteins are unleashed only after the translocon is fully assembled (27–29). Triggering effector secretion requires a conformational change in the translocation pore that, in the case of *Pseudomonas*, is transmitted via the C-terminus of PopD to the collar domain of PcrV (27, 30). The nature of the host cell process that triggers the conformational change in the translocon remains enigmatic.

In this study, we took a genetic approach to map interactions between translocon proteins. We exploited the incompatibilities between the homologous T3SS systems of *P. aeruginosa* and *Aeromonas hydrophila* to map an interaction between the needle tip protein and two small regions of PopB. Functional assays of our genetic hybrids revealed that this interaction is crucial for the formation of the translocation pore, but not for insertion of the translocators into the host membrane. We therefore propose for the first time that the needle tip protein is actively involved in the oligomerization of the translocator proteins into the translocation pore.

## Results

### *P. aeruginosa* and *A. hydrophila* translocator proteins are incompatible

Translocator proteins from *P. aeruginosa* are not interchangeable with their homologs from *Yersinia* spp. (23, 27, 31). We had previously used this observation to genetically map the protein-protein contacts that were disrupted by combining *Pseudomonas* and *Yersinia* translocator proteins. Subsequently, we used hybrid proteins that break a specific contact to assign functions to the interaction. This led to the discovery of several new protein-protein interactions, two of which—an interaction between PopD and PcrV, as well as a dimer interaction between PopD monomers—are important for sensing host cell contact. Here we extend this analysis by examining the incompatibility between *P. aeruginosa* and *A. hydrophila* translocator proteins.

In terms of protein sequence, the translocators from *P. aeruginosa* share 40% identity and 62% similarity with those of *A. hydrophila*. (PcrV-AcrV: 34.2% identical, 51.2% similar; PopB-AopB: 45.9% identical, 70.9% similar; PopD-AopD: 39.1% identical, 62.6% similar; see **Fig. S1**.) AcrV has extra bulk in the tip domain compared to PcrV (23). AlphaFold2 models predict that the extra bulk of AcrV is concentrated in two adjacent 25-residue helices connected with a short loop (32). This feature is completely missing from PcrV, although the rest of the tip domain and central helices are folded similarly (**Fig. 1A**).

**Figure 1.**
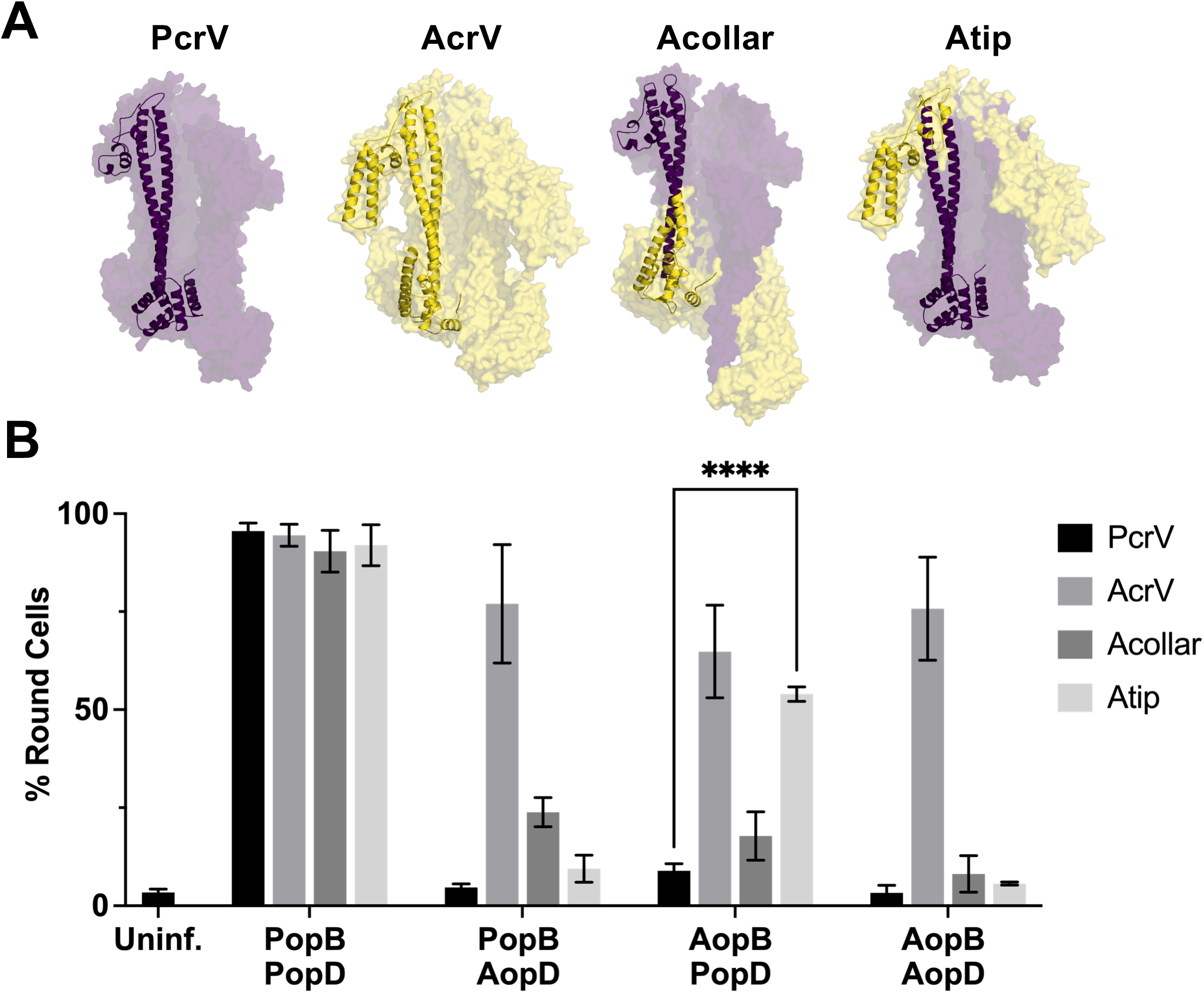
Translocator proteins from *P. aeruginosa* and *A. hydrophila* are not interchangeable. **A.** Models of the needle tip protein from *P. aeruginosa* (PcrV), *A. hydrophila* AH-3 (AcrV), and two chimeric fusion V-proteins (“Atip” and “Acollar”). Our chimeric construct “Acollar” has residues 1-F143 from AcrV fused to PcrV(Q124-end). Our chimeric construct “Atip” has residues 1-A158 from PcrV, A179-S310 from AcrV, and V244-end from PcrV. Monomer models were created from protein sequences using AlphaFold2 implemented in Google Collab Notebooks (32, 54). The position of the N-terminal globular domain (collar domain) assumes various positions in different AlphaFold2 models. The tip domain has a high degree of confidence and is static between models. Pentamer models were created by aligning monomer models with the *Shigella* IpaD pentamer structure (PDB-ID 7rye) (55). Color coding: purple = *Pseudomonas*; yellow = *Aeromonas*. **B**. All strains lack the pore-forming translocator operon (Δ*pcrHpopBD*) and produce the indicated needle tip protein: PcrV (RP3624), AcrV (RP7595), Acollar (RP6166), Atip (RP6425). These background strains were complemented with plasmids carrying the indicated translocator proteins along with both export chaperone homologs *pcrH* and *acrH* (see Table S1 and Table S2). A549 epithelial cells were infected and cellular rounding (a proxy for delivery of ExoS) was monitored by microscopy. ExoS causes actin depolymerization leading to cellular rounding. Statistical significance was calculated by two-way ANOVA with Tukey multiple comparisons test. n=3 biological replicates, SD error bars.

To determine whether translocator proteins from *A. hydrophila* can substitute for their respective *P. aeruginosa* homolog, we constructed *P. aeruginosa* strains carrying each combination of translocator proteins [needle tip (*Pseudomonas* PcrV or *Aeromonas* AcrV), B-translocator (*Pseudomonas* PopB or *Aeromonas* AopB), and D-translocator (*Pseudomonas* PopD or *Aeromonas* AopD)]. Moreover, since we had previously found an interaction of the collar domain of PcrV with PopD, we expanded this analysis to encompass hybrid needle tip proteins in which either the N-terminal collar domain or the tip domain of PcrV had been replaced with the corresponding region of AcrV (**Fig. 1A**). These strains were assayed for their ability to intoxicate epithelial cells (**Fig. 1B** and **S2**). Compatibility among the translocator proteins leads to cell rounding because it permits delivery of the effector ExoS, which causes actin depolymerization. Incompatible systems induce cell rounding more slowly, or not at all (**Fig. S2b**). The native *Pseudomonas* translocators (PopB and PopD) are compatible with both PcrV and AcrV. In contrast, *Aeromonas* AopB and AopD each displayed specific incompatibilities. For example, AopD is only compatible with AcrV. That is, the two systems containing both AopD and AcrV induce ~75% cell rounding within two hours, whereas the strain producing AopD and PcrV induces <25% cell rounding. AopB is similarly incompatible with PcrV, but functional when produced in the context of its cognate needle tip protein, AcrV.

The PcrV-AcrV hybrid “Atip” contains the bulkier tip domain of AcrV substituted for the tip domain of PcrV, while the hybrid “Acollar” has an AcrV substitution in the N-terminal globular domain. Notably, AopB is incompatible with PcrV and Acollar, but significantly rescued by AcrV or Atip. This finding argues for an interaction between the B-translocator and the tip domain of the needle tip protein. Interestingly, AopD function is not fully restored by either hybrid needle tip, arguing that AopD breaks interactions with both the collar and tip domains of PcrV.

The cytotoxicity results shown here confirm that *P. aeruginosa* and *A. hydrophila* pore-forming translocator proteins cannot be freely exchanged. Importantly, the performance of AopB is significantly rescued in the context of chimeric “Atip” compared to wildtype PcrV. We decided to pursue this interaction further since no function has been assigned to the tip domain of the needle-tip protein for any T3SS.

### The tip domain of the needle tip protein interacts with PopB

The tip-dependent phenotype seen in Figure 1 is important because it suggests that PopB interacts with the tip domain of PcrV. Next, we sought to narrow down the specific region of AopB that clashes with the PcrV tip domain. We used an iterative approach to create a panel of PopB-AopB chimeric proteins which were assayed in the context of the wildtype PcrV and the chimeric “Atip” (**Fig. 2, S3, S4**). The performance of these chimeric protein sets during in vivo testing gave us insights into the critical regions needed for the interaction between these proteins.

**Figure 2.**
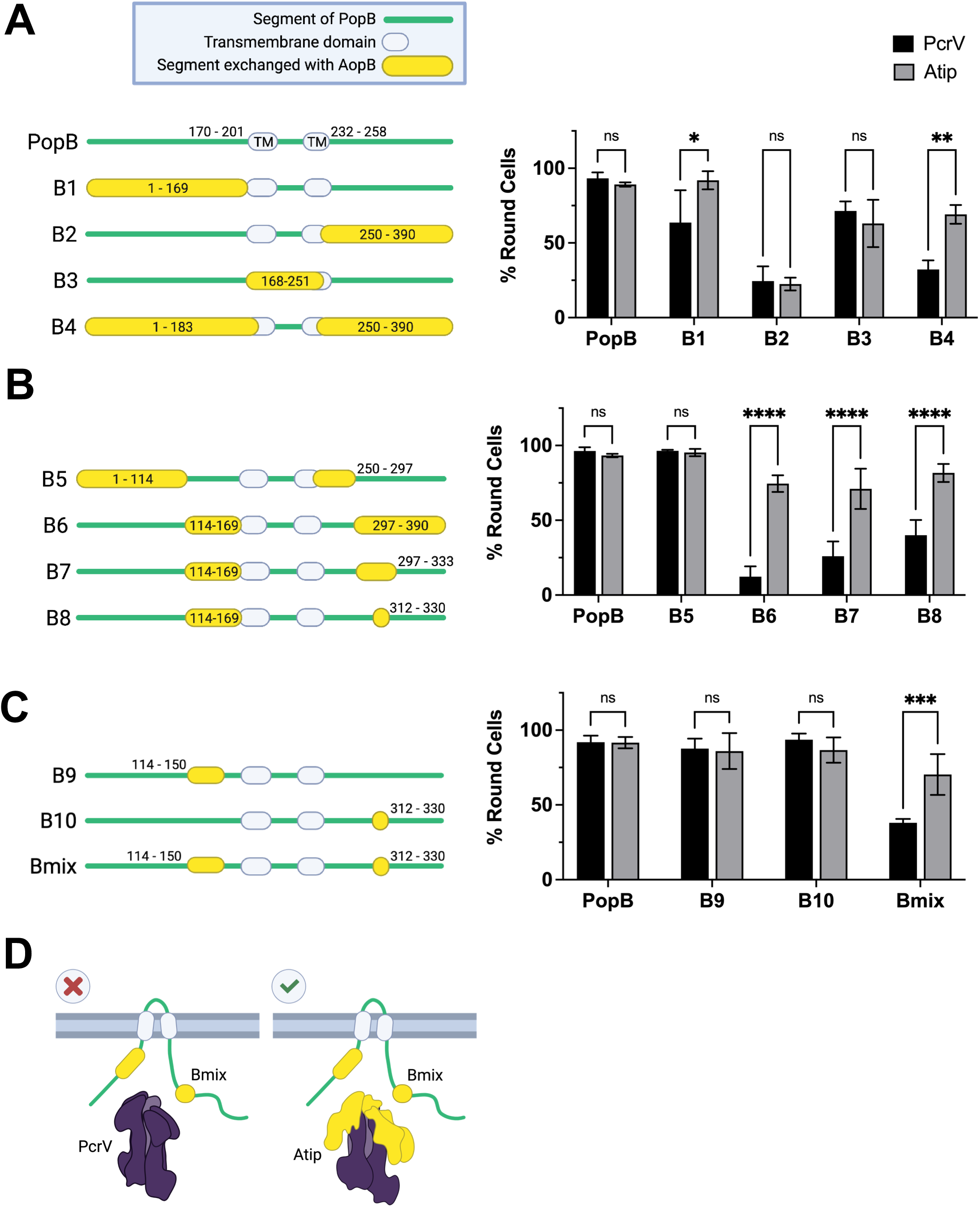
The AopB-Atip interaction is mapped to two small regions of the B-translocator. Chimeric B-translocator proteins were created by fusing together portions of *P. aeruginosa* PopB (purple) and *A. hydrophila* AopB (yellow). The fusion junctions are numbered according to the PopB sequence. Each chimeric B-translocator was tested for cytotoxicity against A549 cells in background strains producing either PcrV (RP3624) or Atip (RP6425). These strains were complemented with plasmids encoding the indicated chimeric B-translocator, *popD*, and both chaperones *pcrH* and *acrH*. Translocation of the effector ExoS into the host cell induces cellular rounding. n=3 biological replicates, SD error bars. Statistical significance calculated using two-way ANOVA with Sidak multiple comparisons test. **A-C. Left:** Schematic diagram of PopB-AopB chimeras. Green line indicates PopB. Yellow bubbles indicate AopB. Residues are numbered according to PopB amino acid sequence. PopB transmembrane domains are 170-201 (TM1) and 232-258 (TM2). Right: Cytotoxicity data for PopB-AopB chimeras. Initial junctions correspond approximately to the two transmembrane domains in PopB.Successively smaller AopB substitutions to narrow down the region of incompatibility. Construct Bmix harbors the minimal region of AopB that still confers tip-dependence. **D.** Schematic diagram of Bmix chimera with wildtype PcrV or chimeric Atip.

PopB contains two transmembrane domains, residues 170-201 and 232-258, and both the N- and C-termini of PopB are in the extracellular space (33). Constructs B1 and B2 that replace the N-terminal and C-terminal extracellular portion of PopB with the corresponding region of AopB resulted in a weak tip-dependent phenotype and loss of function, respectively (**Fig. 2A**). We therefore decided to replace either the central trans-membrane/intracellular domain of PopB (B3) or both extracellular domains (B4). The latter substitution resulted in a chimeric protein that retained the tip-dependent phenotype of AopB. We therefore focused on further narrowing down the portions of the extracellular regions of AopB that clash with the wild-type PcrV needle tip. We subdivided the N- and C-terminal extracellular domains and replaced those regions with the homologous portions of AopB, generating constructs B5 and B6. While B5 was functional in the context of PcrV, construct B6 still required the hybrid Atip needle tip for function. (**Fig. 2B**). Further reduction of the C-terminal substitution (B7, B8) demonstrated that replacing amino acids 312-330 of PopB with the corresponding amino acids from AopB allowed for a construct that maintained the tip-dependent phenotype (**Fig. 2B**). Finally, we were able to reduce the N-terminal substitution to amino acids 114-150 of PopB and still maintain the tip-dependent phenotype (Bmix, **Fig. S3, Fig. 2C**). Notably, neither N- or C-terminal substitution on its own is sufficient to confer the Atip-dependent intoxication phenotype (**Fig. 2C**), arguing that these two regions have to interact to form the interface with the needle-tip. Whether this is an intra-subunit interaction, or whether N- and C-terminus from adjacent PopB proteins interact is unclear **Fig. S4C**).

In the course of mapping the region of incompatibility between AopB and PcrV we noticed that many of the constructs that begin with PopB and end with AopB were inactive (**Fig. S4A**). The converse fusions retained activity (**Fig. S4B**), suggesting that the choice of fusion joint was likely not to be blamed. Indeed sandwich fusions in which the PopB N-terminus is paired with C-terminal portions of PopB restore function (e.g. B4 in **Fig. 2A**), suggesting that the more likely explanation for the lack of function of the constructs is that the mismatch breaks a critical interaction between the N- and C-terminus of PopB. Whether this interaction is intramolecular, or needed for PopB dimer formation (**Fig. S4C**), is unclear.

Finally, we examined whether the added bulk evident in the AcrV needle-tip is required for AopB function. However, deletion of the two 25 amino acid helices unique to AcrV did not result in incompatibility with AopB (Fig. S5), arguing that the incompatibility affects a different, more conserved portion of the PcrV tip domain and that the affected interaction is likely also conserved in the PopB-PcrV pairing.

In summary, we have mapped an interaction between the B-translocator and the needle-tip and generated a construct, Bmix, which breaks a critical contact with PcrV that can be restored by pairing this translocator with the Atip hybrid needle-tip protein. We next exploited these constructs to assign a function to the affected interaction.

### The tip domain of the needle tip protein is not essential for membrane insertion of PopB

We sought to understand which stages of translocon assembly are affected by mismatch between the B-translocator and the needle tip. It has been established that the collar domain of PcrV is required for insertion of PopB and PopD (23). We hypothesized that the tip domain interaction is likewise critical for membrane insertion of PopB and PopD. To examine the effect of breaking the interaction between the PcrV tip domain and the B-translocator, we infected A549 epithelial cells with *P. aeruginosa* strains carrying the chimeric Bmix protein and either wildtype PcrV or chimeric Atip. The amount of translocator proteins in the infection supernatant and in the host cell membrane was assessed by Western blot. There was no significant difference in the amount of membrane-inserted Bmix in the context of wildtype PcrV compared to chimeric Atip, indicating that the tip domain interaction is not in fact required for membrane insertion of PopB (**Fig. 3**).

**Figure 3.**
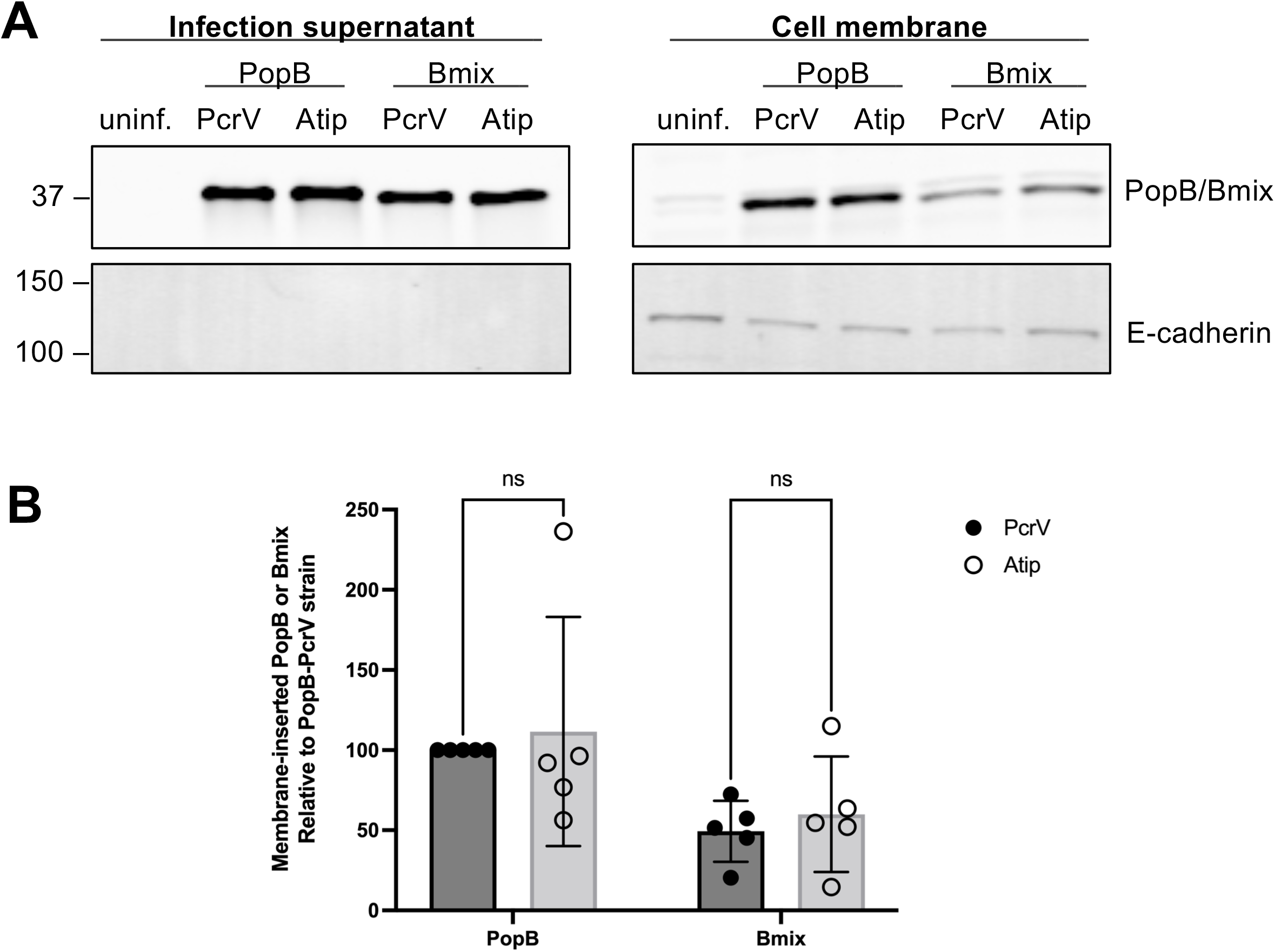
The Bmix-PcrV mismatch does not hinder membrane insertion of PopB. A549 cells were infected for 2h with *P. aeruginosa* producing PopD, as well as the indicated needle-tip/B translocator combination (background strains RP3670 and RP11222). After infection, cells were washed with 1M KCl to remove non-specifically adhered protein, and membrane-inserted translocator proteins were extracted using Triton X-100 (see Methods). **A.** Translocator proteins, as well as E-cadherin (fractionation/loading control) were detected by Western blot in extract and supernatant samples. The blot is representative of five independent experiments. **B.** Quantification of 5 biological replicates. The amount of PopB/Bmix is normalized to E-cadherin or EGFR. SD error bars. Statistical significance was assessed with two-way ANOVA using Sidak multiple comparisons test. Significance threshold: p<0.05.

### Pore formation is hindered by disruption of the interaction between the PopB and the tip domain of PcrV

We next turned our attention to the formation of the translocation pore, and whether pore formation is disabled by disruption of the interface between PopB and PcrV. To answer this question, we measured pore formation in two host cell types. A549 cells were infected in the presence of propidium iodide (PI), which is small enough (668 Daltons) to enter cells through translocation pores (34–36). After two hours of infection, the cells were rinsed and imaged by brightfield and fluorescence microscopy. There was no significant difference in PI uptake between strains producing wildtype PopB and either PcrV or Atip, reiterating our finding from the cytotoxicity assay that PopB can be paired with either version of the needle tip protein (**Fig. 3A-B**). On the other hand, the strain producing Bmix and PcrV averaged less than 25% the amount of PI uptake compared to the strain producing PopB and PcrV, but this defect was significantly corrected in the Bmix-Atip strain. These findings indicate that mismatch between PopB and the tip domain of PcrV specifically impedes the formation of the translocation pore.

In order to corroborate this result, we also tested pore formation in sheep red blood cells, a well-established technique for measuring T3SS pore formation (37). Sheep erythrocytes were pre-treated with papain, which cleaves surface glycoproteins and exposes cryptic binding sites for *P. aeruginosa* hemagglutinin (38). The erythrocytes were infected for one hour, then the amount of hemolysis was measured spectrophotometrically. Data from the hemolysis assay (**Fig. 4C**) closely mirrors that of the propidium iodide uptake assay. Again, there was not a significant difference in the amount of pore formation whether PopB was paired with wildtype PcrV or chimeric Atip. However, Bmix performed significantly better with Atip compared to PcrV, indicating that breaking the interaction between the tip domain of PcrV and PopB is sufficient for abrogating pore formation. These results support the hypothesis that the interaction between PopB and the tip domain of PcrV is essential for oligomerization of the translocation pore.

**Figure 4.**
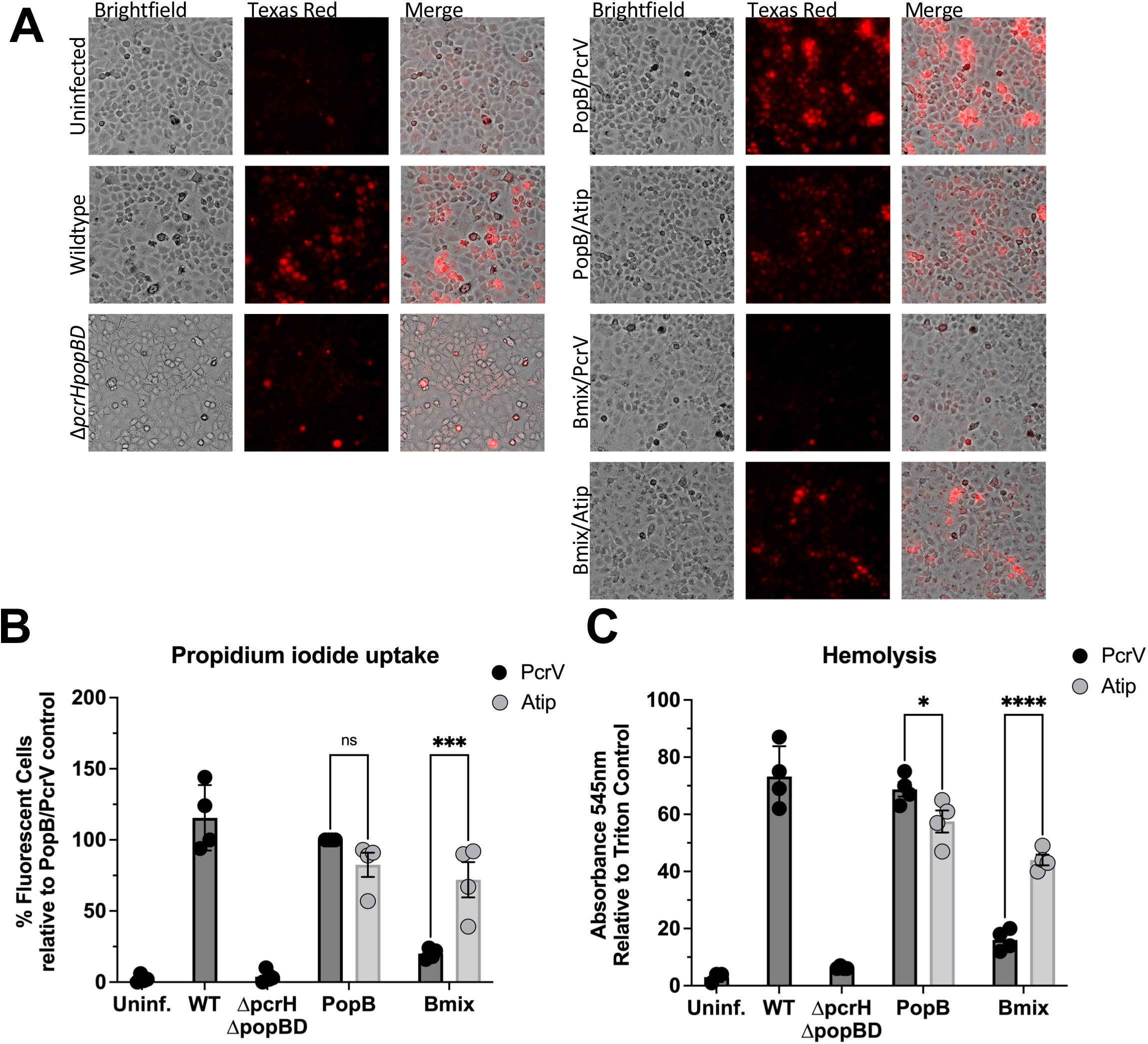
Pore formation is disrupted by mismatch between PopB and PcrV. **A-B.** In the presence of 6 μg/mL propidium iodide, A549 cells were infected with RP11946 (Δ*exoSTY pcrV+* Δ*pcrHpopBpopD*), ”PcrV”, or RP11948 (Δ*exoSTY* Δ*pcrV::pcrV*(*acrVtip*) Δ*pcrHpopBpopD*), “Atip”, producing PopD and the indicated translocators. The wildtype (WT) control strain was RP2317 (Δ*exoSTY pcrV+*). After two hours, cells were washed and imaged (Brightfield and fluorescent with Texas Red filter). **A**. Representative micrographs from one of the biological replicates. **B**. Quantification of 4 biological replicates. Fluorescent cells were scored manually. Each experiment was normalized to the B+V+ strain. Statistical differences analyzed with two-way ANOVA and Sidak multiple comparisons test, * p < 0.05, *** p<0.0005. **C**. Sheep erythrocytes were pre-treated with papain, then infected with RP3670 (*exoS*(G/A-) *pcrV+* Δ*pcrHpopBpopD*), “PcrV”, or RP11222 (*exoS*(G/A-) Δ*pcrV::pcrV(acrVtip)* Δ*pcrHpopBpopD*), “Atip”, producing PopD and the indicated translocators. After two hours, the erythrocyte suspension was mixed, unbroken erythrocytes pelleted, and absorbance of the supernatant measured at 545 nm. Each experiment was normalized to a Triton X-100 lysed control. n=4 biological replicates. Statistical differences analyzed with two-way ANOVA and Sidak multiple comparisons test, * p < 0.05, *** p<0.0005.

### Defective translocation in the mismatched Bmix-PcrV strain was not overcome by bypassing secretion regulation

A hallmark of the T3SS is that effector proteins are secreted only after host cell contact (39, 40). We set out to test whether the mismatch between the PcrV and PopB interrupts the trigger mechanism. The nature of the trigger is unknown but involves a conformational change in the translocon that alters a contact between PopD and the PcrV collar domain (27). In *P. aeruginosa*, the PopN-Pcr1 complex regulates effector secretion before host cell contact (41). Deletion of *pcr1* results in constitutive effector secretion, regardless of host cell contact. While *pcr1*-null strains have dysregulated effector secretion, they are still capable of translocating effectors into host cells in a translocon-dependent manner (27). To test whether the PopB-PcrV interaction is important for secretion regulation, we measured translocation of the effector ExoS in strains lacking the secretion regulator Pcr1. If the PopB-PcrV interaction is primarily required for sensing host-cell contact, deletion of *pcr1* should restore translocation in a strain producing the mismatched Bmix and PcrV. ExoS regulates its own secretion through a feedback loop (42). To remove feedback regulation from the equation and focus solely on translocation, we therefore used strains producing a version of ExoS in which the enzymatic activities have been inactivated by point mutations (ibid.). Translocation of ExoS is impaired by the mismatch between Bmix and PcrV but significantly restored in the Bmix-Atip strain (**Fig. 5**). However, deletion of the secretion regulator *pcr1* is not sufficient to restore ExoS translocation in the mismatched Bmix-PcrV strain, while secretion into the extracellular milieu is not affected (**Fig. S6**). Taken together, these results indicate that the translocation defect of the mismatched Bmix-PcrV strain is primarily due to the defect in pore formation. An additional defect in host cell sensing cannot be ruled out since inhibition of pore assembly could be masking this step.

**Figure 5.**
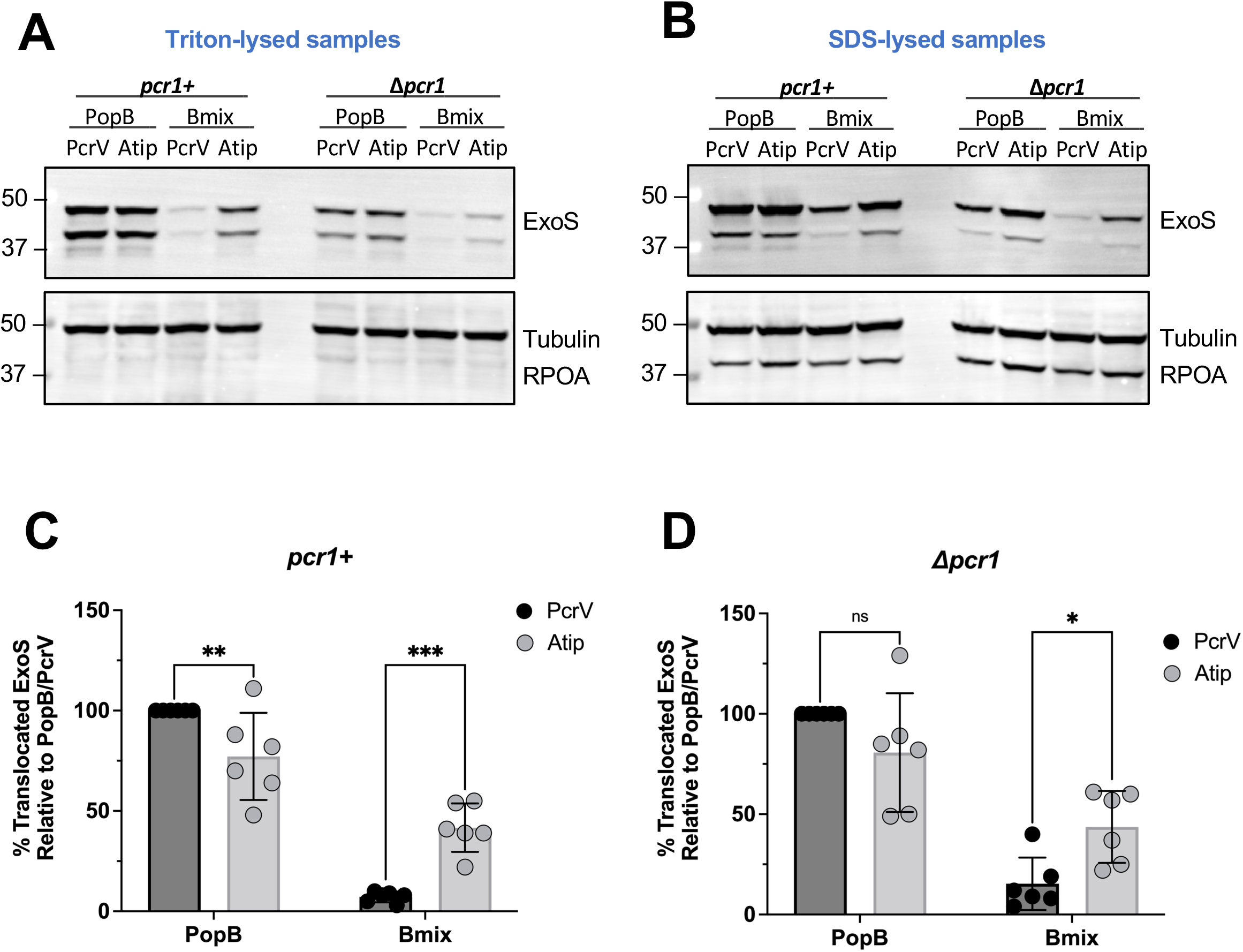
Deletion of the secretion regulator Pcr1 does not restore translocation for the translocator-mismatched (Bmix-PcrV) strain. A549 cells were infected for two hours with strain RP3670 (*exoS*(G/A-) *pcrV+* Δ*pcrHpopBpopD*), ”PcrV”, RP11222 (*exoS*(G/A-) Δ*pcrV::pcrV(acrVtip)* Δ*pcrHpopBpopD*), “Atip”, RP6370 (*exoS*(G/A-) Δ*pcrV::pcrV(acrVtip)* Δ*pcrHpopBpopD* Δ*pcr1*), “PcrV Δ*pcr1*”, or RP11226 (*exoS*(G/A-) Δ*pcrV::pcrV(acrVtip)* Δ*pcrHpopBpopD* Δ*pcr1*), “Atip Δ*pcr1*”, complemented with *popD* and the indicated B-translocator. The amount of translocated ExoS was assessed by lysing the host cells with either Triton X-100, which lyses the eukaryotic cell membrane but does not lyse the bacteria, or treated with SDS, which lyses both the eukaryotic cell and peripherally attached bacteria. ExoS was also monitored in supernatant samples and in bacteria collected from the supernatant after infection. The presence of ExoS, tubulin (host cell cytoplasmic content) and RNA polymerase subunit α (RPOA, bacterial cytoplasmic content) was assessed by Western blot. **A.** The amount of ExoS translocated into A549 cells is shown by the Triton-lysed samples. ExoS is cleaved intracellularly (56). **B.** SDS-lysed samples show the total amount of ExoS both translocated into host cells and remaining in attached bacteria. **C-D.** Quantification of ExoS translocation in Triton-lysed samples. The amount of translocated ExoS is normalized to tubulin and to ExoS in the strain producing PopB/PcrV. n=6 biological replicates. SD error bars. Statistical differences analyzed with two-way ANOVA and Sidak multiple comparisons test, * p<0.05, ** p<0.005, *** p<0.0005.

## Discussion

The T3SS translocon is difficult to study in isolation. By using a genetic approach, we have been able to map protein-protein interactions that are critical for translocon function and use non-functional hybrids to assign functions to these interactions. In this study, we used an incompatibility between translocator proteins of *A. hydrophila* and *P. aeruginosa* to map protein-protein interactions between the needle tip protein and one of the pore-forming translocator proteins. We identified two regions of AopB/PopB that interact, in a concerted manner, with the tip of the needle-tip protein (AcrV/PcrV). Further characterization of the interaction demonstrated that it is not required for secretion or insertion of the pore-forming translocator into the host cell membrane. Rather, the interaction is critical for formation of the translocation pore, which significantly expands the role the needle-tip plays in translocon assembly.

Mapping interactions among translocator proteins has been challenging. Biochemical assays have been used to demonstrate interactions using purified pore-forming translocator proteins (20, 43). Recently, a combination of accessibility of cysteine mutants to chemical modification, as well as disulfide bond formation between adjacent residues, was used to map interactions in the *Shigella flexneri* translocation pore assembled in the plasma membrane of infected cells (44–46). Here we used a genetic approach to identify several interactions that are disrupted by mixing and matching *P. aeruginosa* and *A. hydrophila* translocator proteins (**Fig. 1**). First, our data indicate that AopD fails to interact productively with the *P. aeruginosa* needle-tip protein PcrV. This was not surprising to us, since AopD, just like YopD, harbors a phenylalanine residue at a position corresponding to alanine 292 of PopD. We had shown previously that this substitution is incompatible with *P. aeruginosa* PcrV and substituting the phenylalanine in YopD with alanine (the corresponding amino acid in PopD) largely restored translocator function (27). However, unlike the YopD-PcrV incompatibility, producing AopD in the context of a hybrid needle-tip in which the collar domain of PcrV was replaced with the corresponding region of AcrV, only partially restored function. Since AopD fully supported translocation in the context of AcrV, this result argues that AopD makes two contacts with the needle-tip: one with the collar domain, and a second interaction with the tip of the needle-tip protein. We are currently in the process of mapping this second interaction in order to determine its function in the translocation process. While mapping the incompatibility between AopB and PcrV, we discovered that there is an internal interaction between the N- and C-terminus of PopB that is also disrupted in PopB-AopB hybrids (**Fig. S4**). Function could be restored by having both the N-terminus and C-terminus of the hybrid protein derive from PopB. The interaction that is disrupted in the PopB-AopB hybrids could be intramolecular, i.e. required for the folding of individual PopB monomers, or intermolecular, i.e. required for PopB oligomerization. Further research will be needed to distinguish between these possibilities. Finally, our genetic approach indicates that the very tip of the needle-tip protein interacts with two regions of the B-translocator that both face towards the extracellular milieu (33). The needle-tip dependent translocation defect was only evident when both regions of PopB (residues 114-150 and 312-330) were substituted with the corresponding regions of AopB (resulting in construct Bmix, Fig. 2), suggesting that the two regions both interact with the needle-tip and that a partial substitution is not divergent enough to block the contact with the P. aeruginosa needle-tip protein PcrV. As with the internal PopB interaction, the two interacting regions we identified here could be involved in an intermolecular interaction that allows a PopB monomer to interact with the needle-tip. However, since we know that PopB can dimerize (27), the two regions could also be part of an epitope that is formed by two adjacent PopB monomers, which is then recognized by the needle-tip. A better structural understanding of the translocon will be necessary to resolve this question.

Antibodies directed against the very tip of PcrV are protective in vivo and have been used in clinical trials to treat *Pseudomonas* infections (47). Since this portion of the needle-tip had not been implicated in a particular translocon function, we decided to delve more deeply into the interaction between PopB and the tip domain of PcrV. Using a construct that specifically disrupts this interaction (Bmix) as well as a hybrid needle-tip that restores translocation in the context of Bmix (Atip), we assessed several aspects of translocon function to pinpoint the role of the interaction in effector translocation. First, we assessed the ability of PcrV to insert PopB or Bmix into the host cell membrane. While we recovered less Bmix than PopB in these experiments, the amount was not affected by the needle tip, demonstrating that the Bmix-PcrV incompatibility does not affect Bmix insertion (**Fig. 3**). We next assessed the ability of *P. aeruginosa* to form pores in membranes by detecting translocon-dependent uptake of propidium iodide. Unlike in the translocon insertion experiments, we observed a defect in pore formation when combining Bmix with PcrV, which was reversed by replacing the wild type needle-tip protein with the PcrV(Atip) hybrid protein (**Fig. 4**). Taken together these data argue that the interaction disrupted by co-producing PcrV and Bmix is required for assembly of the translocation pore. We also assayed downstream steps in translocation by detecting translocated ExoS protein in lysates of infected epithelial cells. While we could recapitulate the tip-dependent translocation phenotype of bacteria producing the Bmix translocator protein, this defect could not be restored by removing a negative regulator of effect secretion, Pcr1, from the system (**Fig. 5**). Restoration of effector injection by removing Pcr1 would have indicated that the Bmix-PcrV mismatch interferes with host cell sensing (deleting *pcr1* obviates the need for a host cell trigger). The defect in pore assembly appears to be the primary block in translocon function incurred by the Bmix-PcrV mismatch. However, an additional role of this interaction in docking of the tip to the translocation pore, or in host cell sensing cannot be ruled out, since a block in an earlier step of the translocation process could mask a defect in these later steps.

Our genetic analysis of translocon function has uncovered a novel step in the translocon assembly process: needle-tip guided formation of the translocation pore. We could envision three different models for this activity (**Fig. 6**). First, the needle-tip could simply serve as a tether for inserted translocator proteins, increasing their local concentration and thereby driving pore formation. Second, the needle-tip could play a more active role in the assembly process, serving as a scaffold to fold and assemble inserted translocator proteins into the nascent pore. Thirdly, the translocator proteins could insert and assemble into a pre-pore, which is opened through the interaction with the needle-tip. The latter is reminiscent of an assembly pathway which was suggested for the *Shigella* T3SS, where it was proposed that IpaB may insert into the plasma membrane as a prepore, based on the observation that an *ipaC* null mutant still retains residual ability to lyse red blood cells (48). Which model is correct, or whether multiple aspects of these models (e.g. tethering and chaperoned folding/assembly) play a role in the assembly of the translocation pore awaits further research. It is also not clear whether PopD similarly relies on the needle-tip for assembly into the translocation pore. Intriguingly, our genetic analysis indicates that AopD also requires the tip-domain of AcrV for function. While we have not mapped this interaction yet, it will be interesting to see if this interaction is also specifically required for pore-formation. Notably, purified PopB and PopD form oligomeric complexes on their own, in vitro, as well as hetero-oligomeric complexes if administered simultaneously. However, PopD-complexes cannot be disrupted through the subsequent addition of PopB (20), arguing that this homomeric complex would likely have to be avoided during assembly on the cell surface. Perhaps needle-tip guided pore assembly avoids this dead-end? Our results also have implications for interpreting in vitro models of translocation pore formation. In those studies, PopB and PopD are used to form pores in lipid vesicles, but in the absence of the needle-tip. If the contribution of the needle-tip is to increase the local concentration of translocator proteins, then this function could likely be overcome by increasing the concentration of translocator proteins in those in vitro reactions. However, if the needle-tip serves as a scaffold that helps organize the pore, then this raises the question of how faithfully in vitro assembled translocation pores recapitulate pores formed by infecting bacteria in host cells. Clearly, assembling this key interface between the host cell and the bacterium is more complex than initially anticipated.

**Figure 6.**
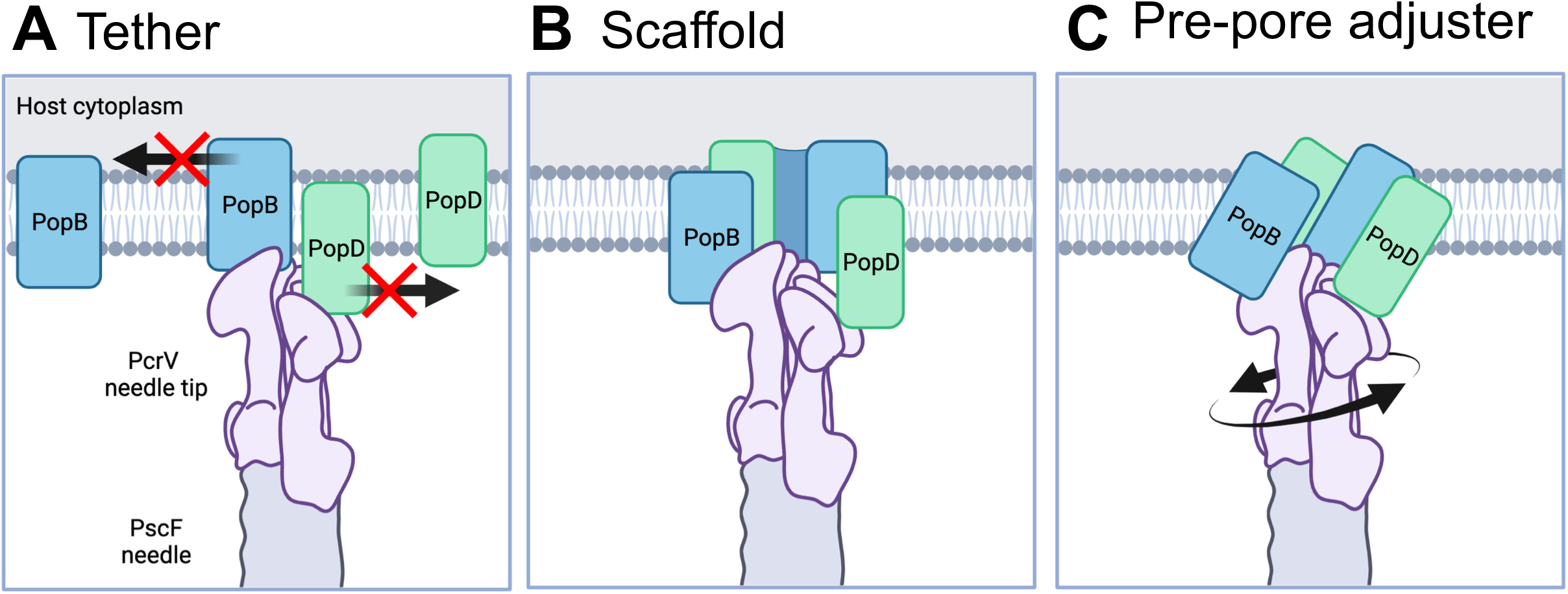
Models of PcrV-assisted translocation pore formation. **A.** PcrV prevents the lateral diffusion of PopB and PopD in the membrane, encouraging oligomerization. **B.** The pore proteins oligomerize around the PcrV needle-tip as they are secreted. **C.** PopB and PopD form a pre-pore structure which is converted by PcrV to its final form. Schematic created with BioRender.com

## Methods

### Bacterial strains, cells, and growth conditions

*E. coli* strains were grown in LB media (10 g tryptone, 5 g yeast extract, 2.5 g NaCl per liter). *P. aeruginosa* strains were grown in “high salt” LB (10 g tryptone, 5 g yeast extract, 11.7 g NaCl per liter with 5 mM MgCl_2_ and 0.5 mM CaCl_2_ and supplemented with 15 μg/mL gentamicin when necessary to retain plasmids. Expression of plasmid-borne genes under control of the *lacUV5* promoter was induced with 100 μM isopropyl β-D-1-thiogalactopyranoside (IPTG).

A549 cells (American Type Culture Collection, Cat. #CCL-185) were grown in RPMI1640 supplemented with 10% FBS (RP10) at 37°C in a 5% CO_2_ atmosphere. Cells were maintained with penicillin and streptomycin. Before infection, A549 cells were rinsed once with Dulbecco’s phosphate buffered saline (DPBS), and the medium was exchanged with RP10 without antibiotics (RP10(-)).

### Plasmid and strain construction

The parent strain was PAO1 (gift from Alain Filloux, (49)). Purified DNA from *Aeromonas hydrophila* AH-3 ((50) was used as a template for *acrH, aopB, aopD*, and *acrV*. Plasmids were constructed using standard molecular biology techniques. Wildtype and chimeric translocator genes were cloned by splicing by overlap extension (SOE) PCR using the primers listed in Table S3. PCR products were digested with the restriction enzymes indicated in **Table S2** and ligated into plasmids pEXG2 (allelic exchange vector), pPSV37 (plasmid that can replicate in *P. aeruginosa* with *lacUV5* promoter and *lacIq*), or pPGEH (plasmid that can replicate in *P. aeruginosa* with T3SS *pcrG* promoter). Mutations in the *P. aeruginosa* chromosome were introduced by allelic exchange as described previously (51). The complete list of strains used in this study is given in Table S1. pPG-*popD* was generated by amplifying three fragments from pP37-*popD (52*) using primers dblatoOri/pGtoRepA, dblatoGent/dlacItoGent, and dlacItoTerm/pGtoPopD which were then combined by Gibson assembly (53), thereby removing the *lacI* gene and remnants of the *bla* ORF in pPSV37, as well as replacing the lacUV5 promoter with the promoter upstream of *pcrG*, and introducing a BbvCI site between the gentamicin resistance gene and replication origin. pPGEH was generated by amplifying the vector backbone of pPG-*popD* using primers pGpolyFor and pGpolyRev and recircularizing the resultant PCR product using GIbson assembly.

### Cytotoxicity assay

Cytotoxicity assays were performed as described previously (27). Briefly, A549 cells were seeded in a 24-well plate at a density of 7.5 × 10^4^ cells/well. The next day, A549 cells were rinsed twice with DPBS and the media was replaced with RP10(-) supplemented with 100 μM IPTG. Bacteria cultures were grown to mid-log (OD_600_ 0.3 - 0.6) in HSLB supplemented with 100 μM IPTG. A549 cells were infected at a multiplicity of infection (MOI) of 25. The infection was stopped by fixation with formaldehyde after 2 hours, except for the kinetics experiments which were stopped at various time points up to 3 hours. Round vs flat (“healthy”) cells were counted manually by low-power phase contrast microscopy.

### A549 membrane insertion assay

A549 membrane insertion assays were performed as described previously (27). A549 were infected for two hours as described for the cytotoxicity assay except for two changes. First, the format is larger (75 cm^2^, tissue culture treated flask). Second, the background strain (RP3670 or RP11222) contains point mutations to abrogate feedback inhibition by ExoS (34, 42). After the infection, a sample of supernatant was collected and the bacteria removed by centrifugation. The A549 cells were rinsed with a high-salt buffer (PBS supplemented with 5 mM MgCl_2_, 0.5 mM CaCl_2_ and 1 M KCl) to break ionic interactions holding peripherally-associated proteins to the membrane. To release intracellular proteins, the A549 cells were treated with the pore-forming toxin streptolysin O (SLO) pre-activated for 20 minutes at room temperature with 10 mM dithiothreitol. Finally, the cells are harvested by scraping and treated with the detergent Triton X-100 to lyse the A549 membrane. Cellular debris was removed by centrifugation, and the solubilized membrane was analyzed by Western blot. Blots were probed for PopB, PopD, and the membrane protein E-cadherin or EGFR was used as a loading control.

### Propidium iodide uptake assay

The day prior to infection, A549 cells were seeded at 9 × 10^5^ cells per well in a 24-well plate. The day of infection, the media was changed to RPMI 1640 without phenol red supplemented with 6 μg/mL propidium iodide (Biotium). Cells were infected at an MOI of 100 for 2 hours, then rinsed twice with PBS-MC. The unfixed cells were imaged with a Cytation 5 imager (BioTek) using a 20X objective for brightfield and Texas Red fluorescent imaging. The number of red cells per field was manually counted.

### Hemolysis assay

Following the previously described protocol (37), sheep erythrocytes in sodium citrate buffer (QuadFive) were washed several times with PBS until the supernatant was clear, then treated with 0.1% w/v papain and 0.01% w/v cysteine for 30 minutes to promote adhesion by *P. aeruginosa* (38). Erythrocytes were washed again and resuspended in RPMI 1640 (without phenol red) to a concentration of 5×10^8^ cells/mL. *P. aeruginosa* strains at mid-log were pelleted gently and resuspended in PBS-MC, the OD_600_ was measured and bacteria were resuspended to a concentration of 2.5 × 10^9^ cells/mL. Bacteria and red blood cells were mixed thoroughly 1:1 (MOI of 5) in a 1.5 mL centrifuge tube. 100 μM IPTG was included in the infection mixture to induce expression of plasmid-borne proteins in *P. aeruginosa*. Samples were centrifuged at 2000x g for 5 minutes to bring the bacteria and erythrocytes into contact, then incubated for 2 hours at 37°C. The samples were resuspended and unbroken cells were removed by centrifugation. Absorbance at 545 nm was read with a BioTek Synergy HT plate reader. The absorbance reading for a PBS blank is subtracted from each reading and samples are normalized to a Triton-lysed sample as follows:

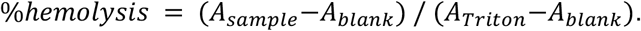

### Translocation assay

The day before the experiment, A549 cells were seeded in 10 cm^2^ tissue culture-treated dishes at 1.5 × 10^6^ cells/dish. A549 epithelial cells were infected with mid-log *P. aeruginosa* for 2 hours at an MOI of 25. The A549 cells were then washed three times with PBS-MC, and rinsed with 1 mL of 250 μg/mL proteinase K in PBS-MC. The protease solution was removed promptly, and the cells were incubated at room temperature for 15 minutes to digest extracellular protein. The protease-treated cells are resuspended in 1ml of PBS-MC with 2 mM PMSF then and pelleted gently (3 minutes, 8000x g). The cells were resuspended in 95μL of PBS-MC with freshly prepared 0.1% Triton X-100 and incubated on ice for 15 minutes. 45μL of the cell suspension is removed and mixed with 15μl of 4x SDS sample buffer (SDS sample). The remaining cells were pelleted, and 45μL of supernatant was removed and combined with 15μL of 4x SDS sample buffer (Triton solubilized fraction). The samples were heated for 10 min at 95°C, separated by SDS-PAGE and analyzed by Western blot. Membranes were probed with antibodies directed against ExoS, tubulin, and RNA polymerase α (RpoA).

### Western blotting

Samples in SDS buffer are heated at 95°C for 15 minutes. Unless otherwise noted, samples are separated on 10% SDS-PAGE gels (BioRad) and transferred to Immobilon-FL PVDF (for fluorescent imaging) or Immobilon-PSQ PVDF (for chemiluminescent imaging). For fluorescent detection, blots were incubated with LI-COR anti-mouse 700 nm or anti-rabbit 800nm secondary antibodies at 1:10,000 concentration and imaged with a LI-COR Odyssey system. Densitometry analysis was performed with LI-COR Odyssey software and ImageJ. For chemiluminescent detection, blots were incubated with goat anti-rabbit or goat anti-mouse secondary antibodies conjugated to horseradish peroxidase and detection was performed with chemiluminescent substrate (Advansta Sirius) with image detection on a GE ImageQuant Las4000 imager.

### Statistical analysis

Statistical analysis was performed in GraphPad Prism using at least three biological replicates performed on independent days using independent bacterial cultures. Statistical tests used are indicated in the figure legends.

## Supporting information

combined supplementary data

## Acknowledgements

This work was supported by NIH grants R21 AI107131 and 5R01EY022052. ECK was supported in part by NIH grant T32 GM007250. Bacterial strain *P. aeruginosa* PAO1F was a gift from Dr. Alain Filloux (Imperial College London). Bacterial strain *Aeromonas hydrophila* AH-3 was a gift from Dr. Juan Tomas (Universitat de Barcelona, Department de Microbiologia). The anti-EGFR antibody was a gift from Dr. Cathleen Carlin (Case Western Reserve University). We are grateful to Dr. George Dubyak (Case Western Reserve University) for the use of the BioTek Cytation 5 plate reader.

## Author Contributions

Conceived and designed the experiments: AR. Performed the experiments: ECK, JDA, JT, AR. Analyzed the data: ECK, AR. Wrote the paper: ECK, AR.

