## Supplementary material for "PopB-PcrV Interactions are Essential for Pore Formation in the *Pseudomonas aeruginosa* Type III Secretion System Translocon": combined supplementary data

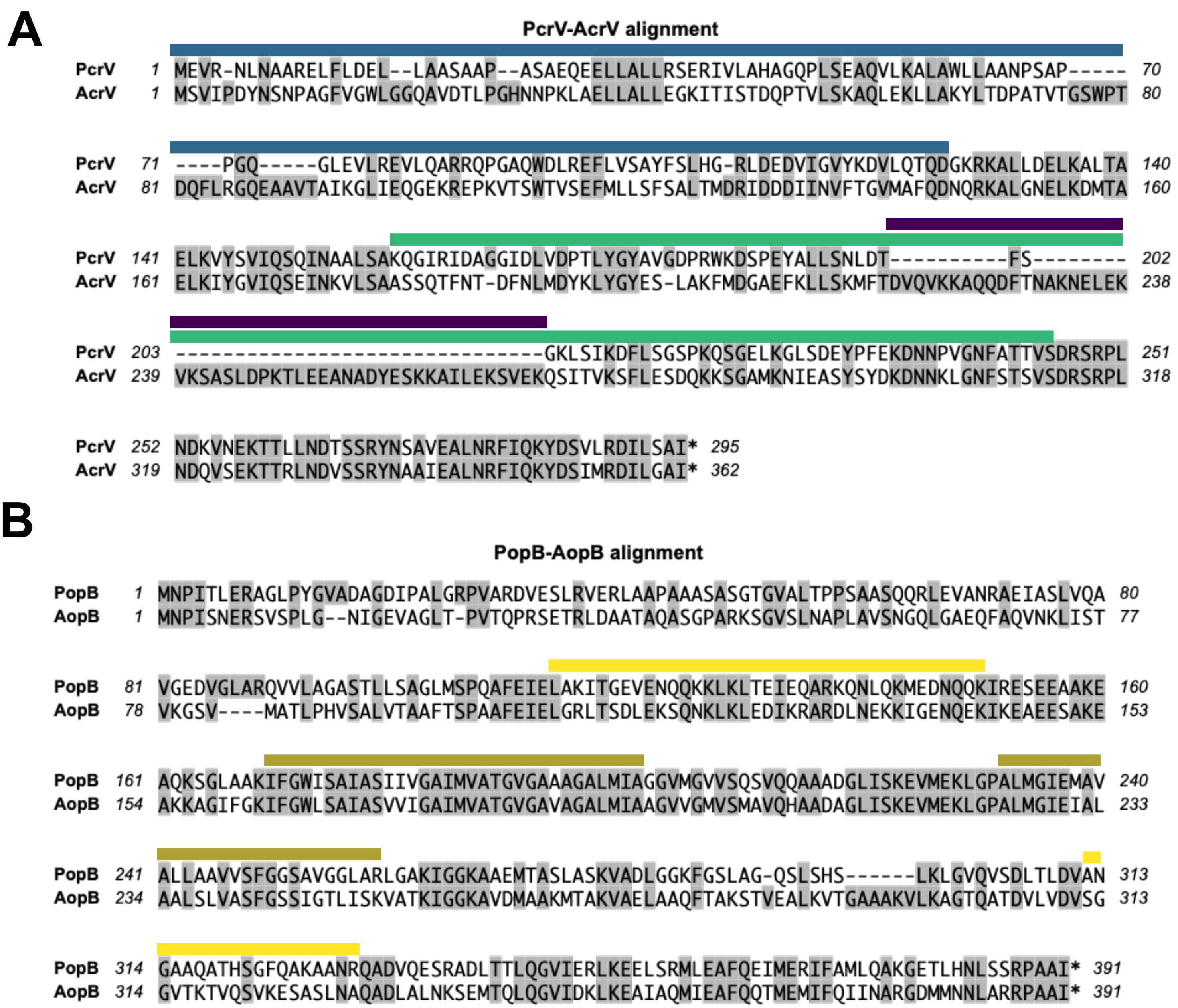

**Figure S1. Protein sequence alignments for PopB-AopB and PcrV-AcrV.** ***Related to Figures 1 and 2.***

**A.** PcrV (Genbank accession AE004091) and AcrV (Genbank accession AY528667) were aligned using ClustalW. The amino acid sequences are 36% identical (50% similar). The exchanged region for the Acollar chimera is indicated above the sequence in blue bars. The exchanged region for the Atip chimera is indicated with green bars above the sequence. The segment that was deleted from Atip to form the Atip-trim chimera (Fig. S5) is indicated in purple bars.

**B.** PopB (from Genbank accession AE004091) and AopB (from Genbank accession AY528667) were aligned using ClustalW. The amino acid sequences are 46% identical (71% similar). Transmembrane domains (residues 170-201 and 232-258) are indicated by olive bars above the sequence. Exchanged regions in the chimeric construct Bmix (residues 114-150 and 312-330) are indicated in yellow bars above the sequence.

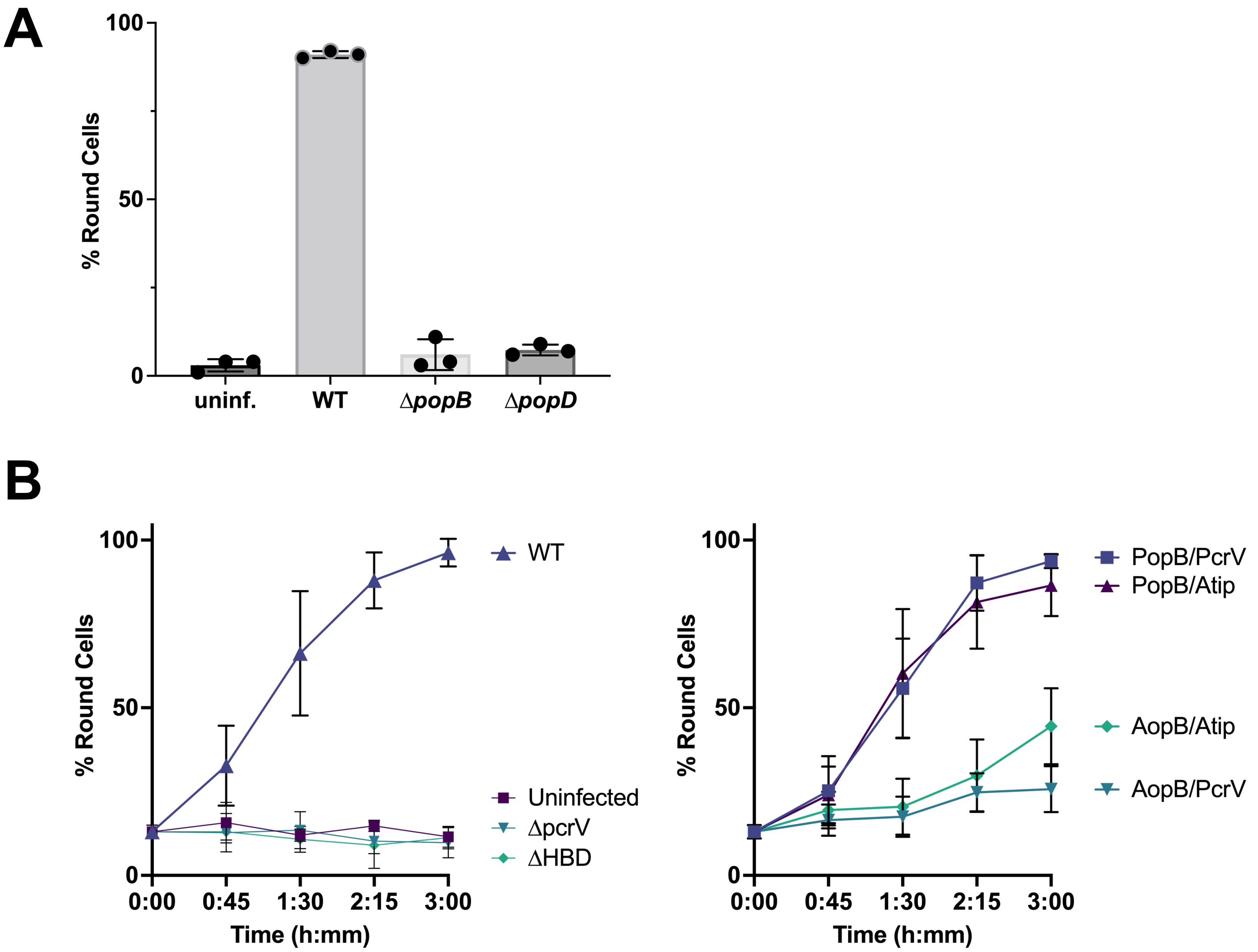

**Figure S2. Additional controls and kinetics for cytotoxicity assay. *Related to Figure 1.***

**A.** The experiment was performed as described in Figure 1 using background strains RP3624 (PcrV) and RP6425 (Atip). The wildtype strain is RP2318. The Δ*pcrV* strain is RP3223. The Δ*popB* and Δ*popD* strains have the background strain RP3624 complemented with a plasmid carrying either *popB* or *popD* along with both export chaperone homologs *pcrH* and *acrH*. n=3 biological replicates. SD error bars.
**B.** Kinetics of cell rounding in response to *P. aeruginosa* infection. The cytotoxicity experiment was performed as described in Figure 1 but stopped at various time points up to 3 hours. n=4 biological replicates. SD error bars. The dataset is split into two graphs for clarity.

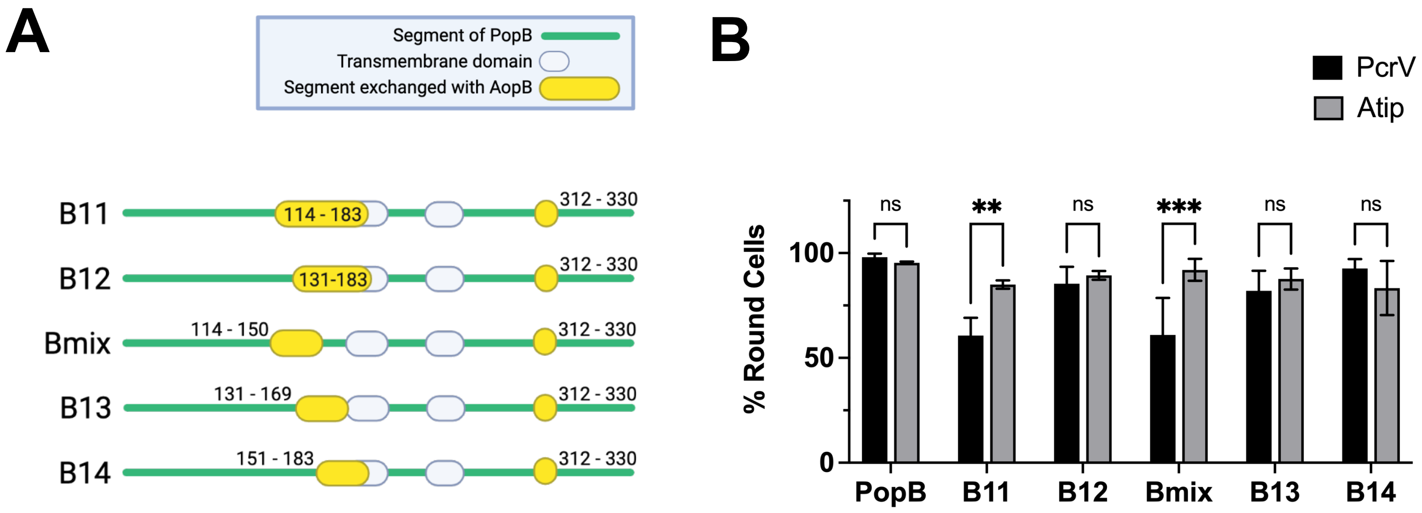

**Figure S3. Refining the region of interest for the PopB-AopB interaction.** ***Related to Figure 2.***

**A.** Schematic diagrams of PopB-AopB chimeras. This set of chimeras was designed to refine the boundaries of the PopB-PcrV interaction, focusing on the N-terminal substitution. Created with BioRender.com

**B**. Cytotoxicity data for PopB-AopB chimeras. The experiment was performed as described in Figure 1 using background strains RP3624 (PcrV) and RP6425 (Atip). n=3 biological replicates. SD error bars. Statistical differences analyzed with two-way ANOVA and Sidak multiple comparisons test, * $p<0.05$, ** $p<0.005$, *** $p<0.0005$, *** $p<0.0001$.

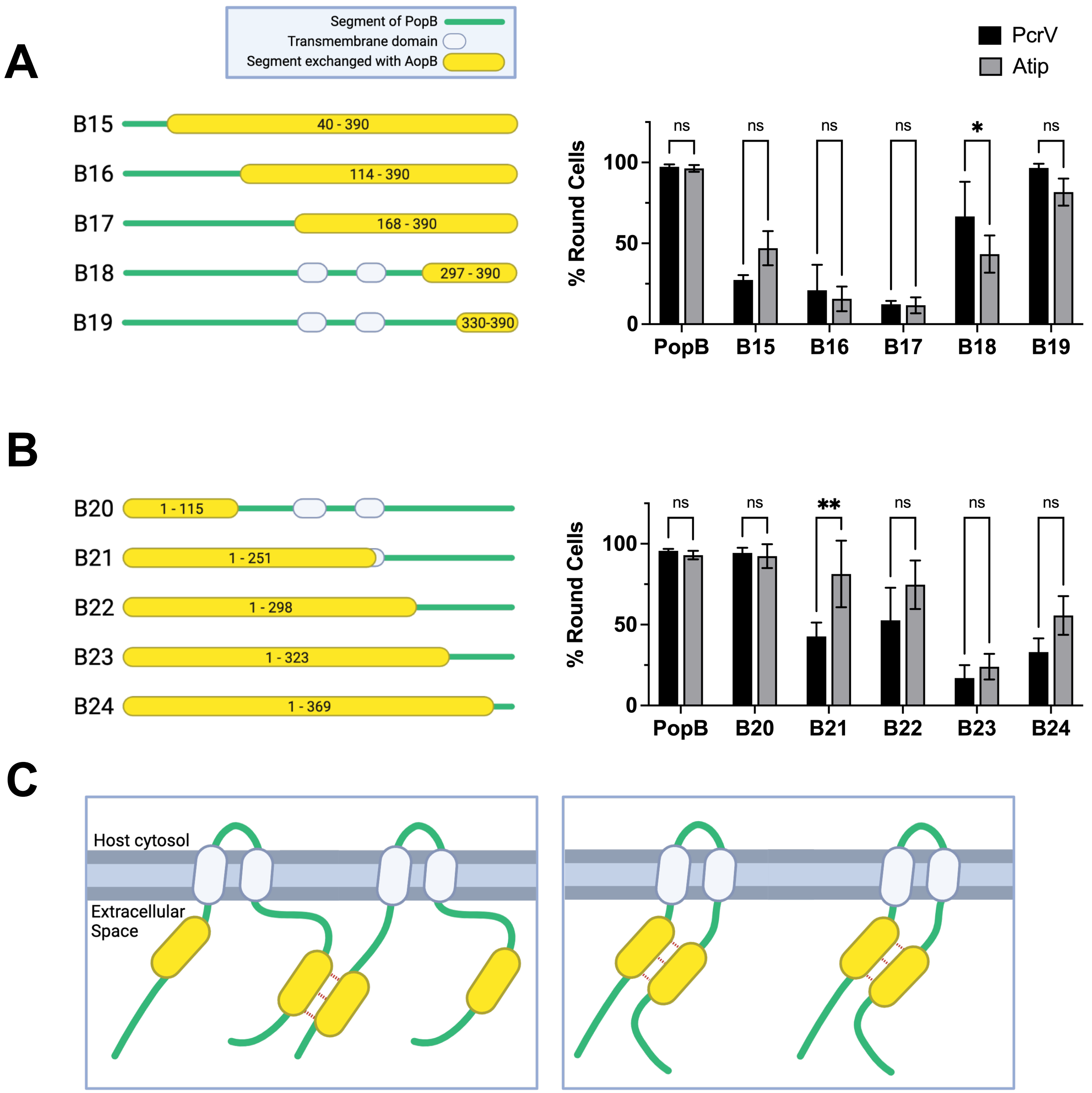

**Figure S4. Two regions of PopB may interact with each other.** ***Related to Figure 2.***

**A-B, left-hand side:** Schematic diagrams of PopB-AopB chimeras.
**A-B, right-hand side:** Cytotoxicity data for PopB-AopB chimeras. The experiment was performed as described in Figure 1 using background strains RP3624 (PcrV) and RP6425 (Atip). n=3 biological replicates. SD error bars. Statistical differences analyzed with two-way ANOVA and Sidak multiple comparisons test, * $p<0.05$, ** $p<0.005$, *** $p<0.0005$, *** $p<0.0001$.
**C**. Schematic diagram showing two possible PopB-PopB interactions. In the first scenario (left-hand panel), two adjacent monomers of PopB interact with each other. In the second scenario (right-hand panel), PopB folds on itself and has an intramolecular interaction. Created with BioRender.com

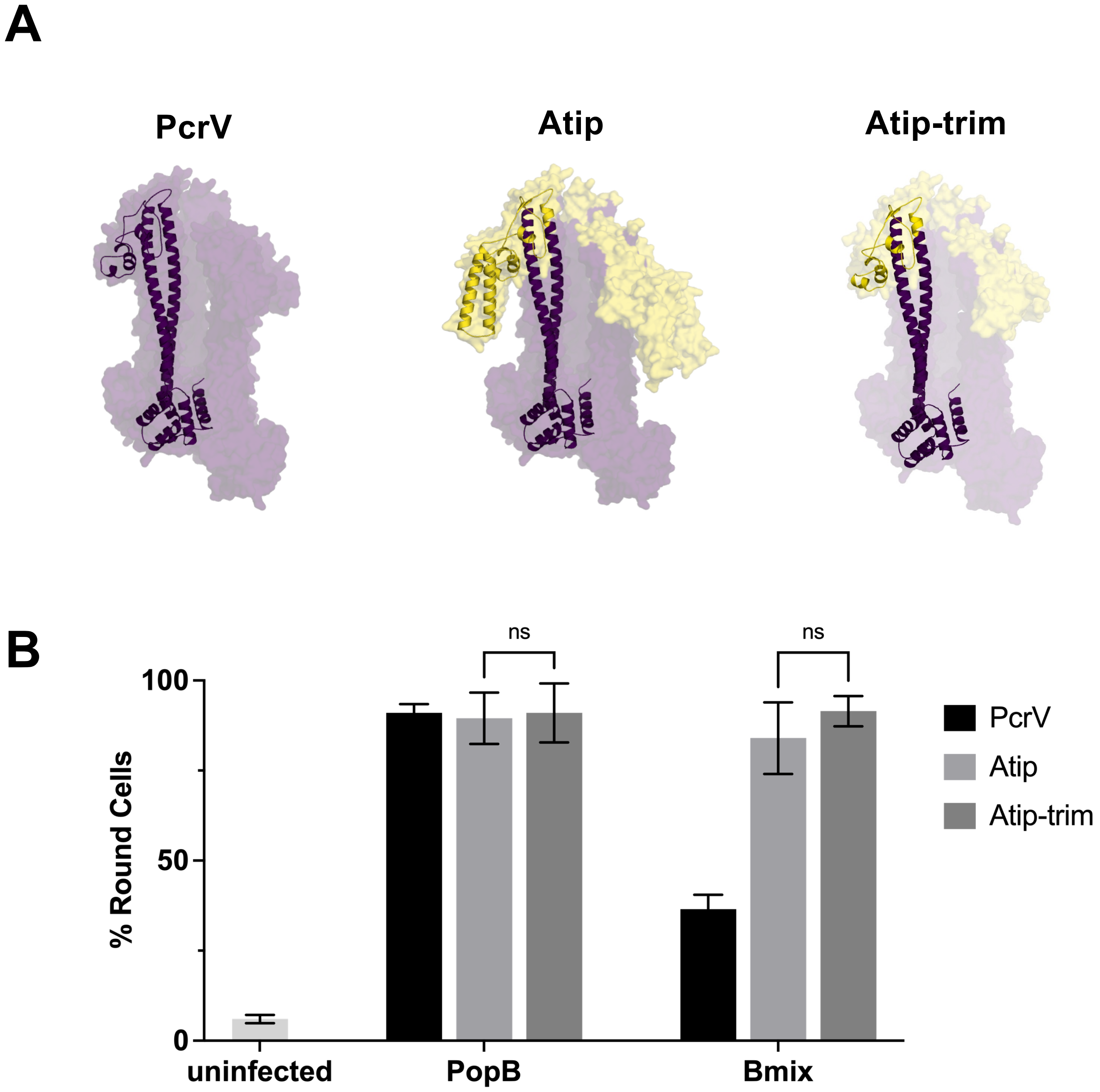

**Figure S5. The non-homologous segment of AcrV is not necessary for rescue of Bmix.** ***Related to Figure 2.***

**A.** Residues 220-269 of AcrV, which have no homologous segment in PcrV, were deleted from the Atip construct to create the PcrV-AcrV chimera "Atip-trim." Models of PcrV, Atip and Atip-trim were created as described in Figure 1.

**B.** The experiment was performed as described in Figure 1 using background strains RP3624 (PcrV), RP6425 (Atip), and RP12517 (Atip-trim). Statistical significance was calculated by two-way ANOVA with Tukey multiple comparisons test. Data bars are a summary of three biological replicates with SD error bars. ns: not significant ($p>0.05$).

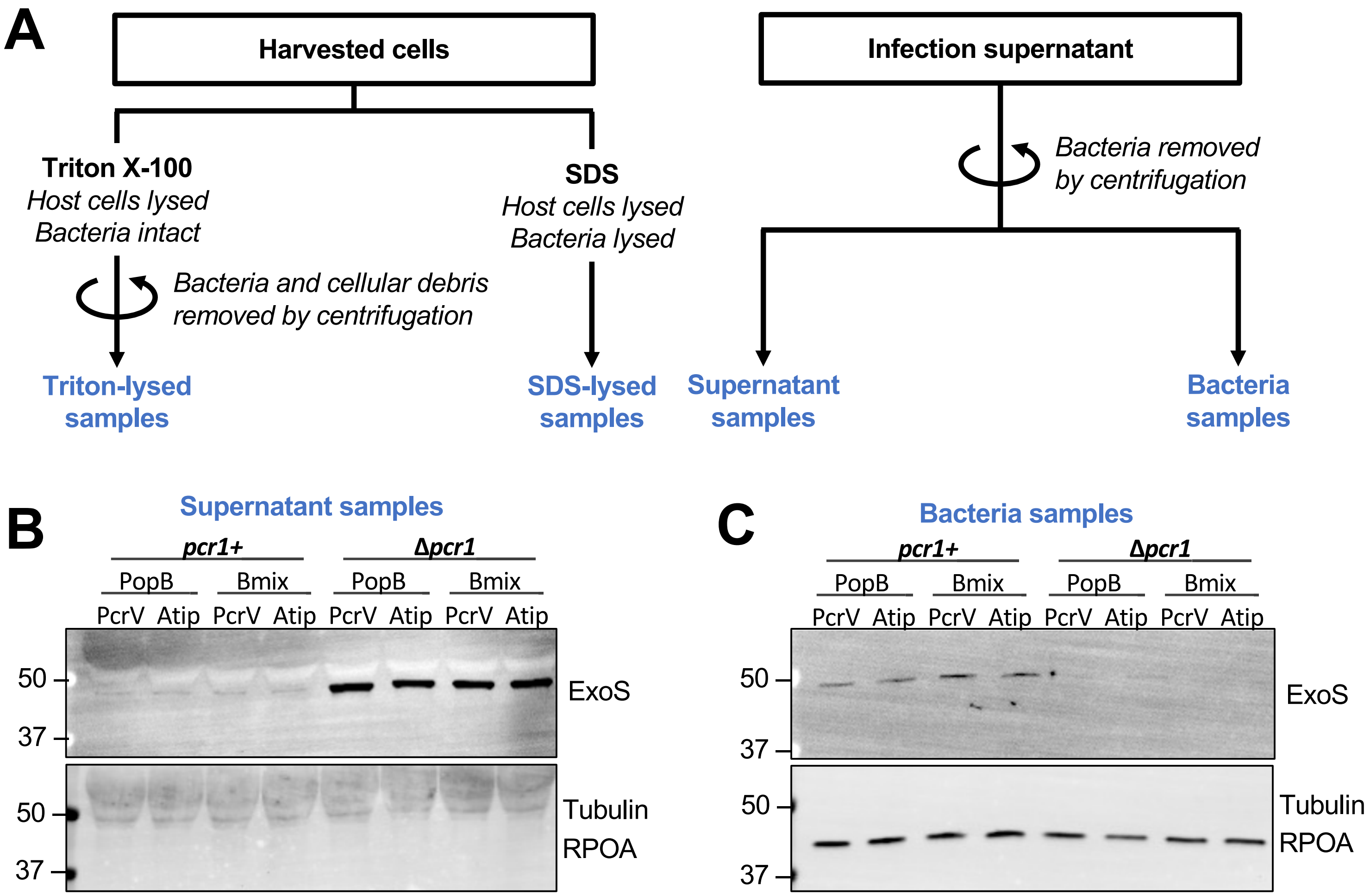

**Figure S6. Supernatant samples for the translocation assay.** ***Related to Figure 5.***

The experiment was performed as described in Figure 5 using background strains RP3670 (PcrV), RP11222 (Atip), RP6370 (PcrV Δ*pcr1*) or RP11226 (Atip Δ*pcr1*).

**A.** Schematic diagram of experimental design.

**B.** Supernatant samples show the amount of ExoS secreted before host-cell contact. Deletion of the secretion regulator *pcr1* causes premature secretion of ExoS.

**C.** Bacteria samples show the amount of ExoS contained in the bacteria cytosol at the end of the infection.

Tubulin (eukaryotic cellular protein) and RpoA (RNA polymerase alpha subunit) serve as fractionation controls.

| **Table S1: Strains** | | | |
|---|---|---|---|
| **Strain #** | **Genotype** | **Ref.** | **Figure** |
| RP2318 | PAO1F Δ*exsE* Δ*exoT* Δ*exoY* | Cisz 2008 | Fig S1 |
| RP3624 | PAO1F Δ*exsE* Δ*exoT* Δ*exoY* Δ*pcrHpopBD pcrV+* | Armentrout 2016 | Fig 1, 2, S1, S3, S4, S5 |
| RP6425 | PAO1F Δ*exsE* Δ*exoT* Δ*exoY* Δ*pcrHpopBD* Δ*pcrV::pcrV(acrVtip)* | This study | Fig 1, 2, S1, S3, S4, S5 |
| RP6166 | PAO1F Δ*exsE* Δ*exoT* Δ*exoY* Δ*pcrHpopBD* Δ*pcrV::acrV(N)-pcrV(C)* | This study | Fig 1 |
| RP7595 | PAO1F Δ*exsE* Δ*exoT* Δ*exoY* Δ*pcrHpopBD* Δ*pcrGV::acrGV* | This study | Fig 1 |
| RP3223 | PAO1F Δ*exsE* Δ*exoT* Δ*exoY “S+”* Δ*pcrV2* | This study | Fig S1 |
| RP2349 | PAO1F Δ*exsE* Δ*exoT* Δ*exoY exoS (R146K/E379D/E381D) "G/A-"* | Cisz 2008 | |
| RP3670 | PAO1F Δ*exsE* Δ*exoT* Δ*exoY exoS (R146K/E379D/E381D) "G/A-"* Δ*pcrHpopBD pcrV+* | Armentrout 2016 | Fig 3, 4, 5, S6 |
| RP11222 | PAO1F Δ*exsE* Δ*exoT* Δ*exoY exoS (R146K/E379D/E381D) "G/A-"* Δ*pcrHpopBD* Δ*pcrV::pcrV(acrVtip)* | This study | Fig 3, 4, 5, S6 |
| RP6370 | PAO1F Δ*exsE* Δ*exoT* Δ*exoY exoS(GAP-/ADPR-)* Δ*pcrHpopBD pcrV+* | Armentrout 2016 | Fig 5, S6 |
| RP11226 | PAO1F Δ*exsE* Δ*exoT* Δ*exoY exoS(GAP-/ADPR-)* Δ*pcrHpopBD* Δ*pcrV::pcrV(acrVtip)* | This study | Fig 5, S6 |
| RP2317 | PAO1F Δ*exsE* Δ*exoS* Δ*exoT* Δ*exoY* | Cisz 2008 | Fig 4 |
| RP11946 | PAO1F Δ*exsE* Δ*exoS* Δ*exoT* Δ*exoY* Δ*pcrHpopBD pcrV+* | This study | Fig 4 |
| RP11948 | PAO1F Δ*exsE* Δ*exoS* Δ*exoT* Δ*exoY* Δ*pcrHpopBD* Δ*pcrV::pcrV(acrVtip)* | This study | Fig 4 |
| RP12517 | PAO1F Δ*exsE* Δ*exoT* Δ*exoY S+* Δ*pcrHpopBD* Δ*pcrV::pcrV(acrVtip-trim)* | This study | Fig S5 |

**Table S2: Plasmids**

| Hybrid Construct | Full name | Description | Ref. |
|---|---|---|---|
| | pPSV37 | colE1 oriR, gentR, PA origin, oriT, lacUV5 promoter, lacIq, stops in every reading frame preceding the MCS and T7 terminator following the MCS relative to the lacUV5 promoter | Lee 2010 |
| | pEXG2 | Allelic exchange vector, colE1 origin, oriT, gentR, sacB | Rietsch 2005 |
| | pPGEH | pPGEH has the pcrG promoter, gent resistance, BbvCI site. I made it by removing popD from pPG-popD and adding back the pPSV37 polylinker (EcoRI to HindIII) | This study |
| | pEXG2-ΔpcrHΔpopBΔpopD | Allelic exchange vector designed to delete from pcrH codon 5 to popD codon 268 | Tomalka 2012 |
| | pEXG2-pcrV(acrVtip) | Exchange plasmid for pcrV::Atip | This study |
| | pEXG2-acrV(N)-pcrV(C) | Exchange plasmid for pcrV::Acollar | This study |
| | pEXG2-ΔpcrV::acrV | Exchange plasmid for pcrV::AcrV | This study |
| | pEXG2-Δpcr1 | Allelic exchange vector designed to delete codons 10–81 of pcr1 | Tomalka 2012 |
| | pEXG2-Atip-trim | Exchange plasmid for pcrV::Atip-trim | This study |
| | pEXG2-ΔpcrV2 | pEXG2 with pcrV deletion allele (Δ4-180) | Cisz 2008 |
| | pP37-acrHpcrH-popBD | acrH, pcrH, popB and popD under the control of the lacUV5 promoter in pPSV37 | Armentrout 2016 |
| | pP37-acrHpcrH-aopBD | Plasmid encoding acrH, pcrH, aopB and aopD under control of the lacUV5 promoter in pPSV37 | This study |
| | pP37-acrHpcrH-popB-aopD | Plasmid encoding acrH, pcrH, popB and aopD under control of the lacUV5 promoter in pPSV37 | This study |
| | pP37-acrHpcrH-aopBpopD | Plasmid encoding acrH, pcrH, aopB and popD under control of the lacUV5 promoter in pPSV37 | This study |
| | pP37-acrHpcrHpopD | Plasmid encoding acrH, pcrH, and popD under control of the lacUV5 promoter in pPSV37 | This study |
| | pP37-acrHpcrHpopB | Plasmid encoding acrH, pcrH and popB under control of the lacUV5 promoter in pPSV37 | This study |
| B1 | pP37-acrHpcrH-APB 169-popD | Plasmid encoding acrH, pcrH, a aopB(1-161G)-popB(169K-390) hybrid and popD under the control of the lacUV5 promoter in pPSV37 | This study |
| B2 | pP37-acrHpcrH-PAB 250-popD | Plasmid encoding acrH, pcrH, popD and a popB(1-250G)-aopB(244S-390) hybrid under the control of the lacUV5 promoter in pPSV37 | This study |
| B3 | pP37-acrHpcrH-PAPB 168-251-popD | Plasmid encoding acrH, pcrH, popD and a popB(1-168A)-aopB(162K-243G)-popB(251G-390) hybrid under the control of the lacUV5 promoter in pPSV37 | This study |
| B4 | pP37-acrHpcrH-APAB 183-250-popD | Plasmid encoding acrH, pcrH, popD and a aopB(1-175I)-popB(183G-250G)-aopB(244S-390) hybrid under the control of the lacUV5 promoter in pPSV37 | This study |
| B5 | pP37-acrHpcrH-APAPB 115-250-298-popD | Plasmid encoding acrH, pcrH, popD and a aopB(1-107L)-popB(115A-250G)-aopB(244S-297V)-popB(298-390) hybrid under the control of the lacUV5 promoter in pPSV37 | This study |
| B6 | pP37-acrHpcrH-PAPAB 114-169-297-popD | Plasmid encoding acrH, pcrH, popD and a popB(1-114L)-aopB(108G-161G)-popB(169K-296H)-aopB(270A-390) hybrid under the control of the lacUV5 promoter in pPSV37 | This study |
| B7 | pP37-acrHpcrH-PAPAPB 114-169-297-333-popD | Plasmid encoding acrH, pcrH, popD and a popB(1-114L)-aopB(108G-161G)-popB(169K-296H)-aopB(270A-332A)-popB(333D-390) hybrid under the control of the lacUV5 promoter in pPSV37 | This study |
| B8 | pP37-acrHpcrH-PAPAPB 114-169-312-330-popD | Plasmid encoding acrH, pcrH, popD and a popB(1-114L)-aopB(108G-161G)-popB(169K-311V)-aopB(312S-332A)-popB(333D-390) hybrid under the control of the lacUV5 promoter in pPSV37 | This study |
| B9 | pP37-acrHpcrH-PAPB 114-150-popD | Plasmid encoding acrH, pcrH, popD and a popB(1-114L)-aopB(108G-143K)-popB(150K-390) hybrid under the control of the lacUV5 promoter in pPSV37 | This study |
| B10 | pP37-acrHpcrH-PAPB 312-330-popD | Plasmid encoding acrH, pcrH, popD and a popB(1-311V)-aopB(312S-330A)-popB(331Q-390) hybrid under the control of the lacUV5 promoter in pPSV37 | This study |
| Bmix | pP37-acrHpcrH-PAPAPB 114-150-312-330-popD | Plasmid encoding acrH, pcrH, popD and a popB(1-114L)-aopB(108G-143K)-popB(150K-311V)-aopB(312S-330A)-popB(331Q-390) hybrid under the control of the lacUV5 promoter in pPSV37 | This study |
| B11 | pP37-acrHpcrH-PAB 40-popD | Plasmid encoding acrH, pcrH, popD and a popB(1-40L)-aopB(38T-390) hybrid under the control of the lacUV5 promoter in pPSV37 | This study |
| B12 | pP37-acrHpcrH-PAPAPB 114-183-312-330-popD | Plasmid encoding acrH, pcrH, popD and a popB(1-114L)-aopB(108G-175I)-popB(183G-311V)-aopB(312S-330A)-popB(331Q-390) hybrid under the control of the lacUV5 promoter in pPSV37 | This study |
| B13 | pP37-acrHpcrH-PAPAPB 131-183-312-330-popD | Plasmid encoding acrH, pcrH, popD and a popB(1-131T)-aopB(126I-175I)-popB(183G-311V)-aopB(312S-330A)-popB(331Q-390) hybrid under the control of the lacUV5 promoter in pPSV37 | This study |
| B14 | pP37-acrHpcrH-PAPAPB 131-169-312-330-popD | Plasmid encoding acrH, pcrH, popD and a popB(1-131T)-aopB(125D-175I)-popB(183G-311V)-aopB(312S-330A)-popB(331Q-390) hybrid under the control of the lacUV5 promoter in pPSV37 | This study |
| B15 | pP37-acrHpcrH-PAPAPB 151-183-312-330-popD | Plasmid encoding acrH, pcrH, popD and a popB(1-151I)-aopB(145K-175I)-popB(183G-311V)-aopB(312S-330A)-popB(331Q-390) hybrid under the control of the lacUV5 promoter in pPSV37 | This study |
| B16 | pP37-acrHpcrH-PAB 114-popD | Plasmid encoding acrH, pcrH, popD and a popB(1-114L)-aopB(108G-390 hybrid under the control of the lacUV5 promoter in pPSV37 | This study |
| B17 | pP37-acrHpcrH-PAB 168-popD | Plasmid encoding acrH, pcrH, popD and a popB(1-168A)-aopB(162K-390) hybrid under the control of the lacUV5 promoter in pPSV37 | This study |
| B18 | pP37-acrHpcrH-PAB 297-popD | Plasmid encoding acrH, pcrH, popD and a popB(1-296H)-aopB(270A-390) hybrid under the control of the lacUV5 promoter in pPSV37 | This study |
| B19 | pP37-acrHpcrH-PAB 330-popD | Plasmid encoding acrH, pcrH, popD and a popB(1-330R)-aopB(331Q-390) hybrid under the control of the lacUV5 promoter in pPSV37 | This study |
| B20 | pP37-acrHpcrH-APB 115-popD | Plasmid encoding acrH, pcrH, popD and a aopB(1-107L)-popB(115A-390) hybrid under the control of the lacUV5 promoter in pPSV37 | This study |

**Table S2: Plasmids**

| Hybrid Construct | Full name | Description | Ref. |
|---|---|---|---|
| B21 | pP37-acrHpcrH-APB 251-popD | Plasmid encoding acrH, pcrH, popD and a aopB(1-243G)-popB(251G-390) hybrid under the control of the lacUV5 promoter in pPSV37 | This study |
| B22 | pP37-acrHpcrH-APB 298-popD | Plasmid encoding acrH, pcrH, popD and a aopB(1-297V)-popB(298L-390) hybrid under the control of the lacUV5 promoter in pPSV37 | This study |
| B23 | pP37-acrHpcrH-APB 323-popD | Plasmid encoding acrH, pcrH, popD and a aopB(1-322V)-popB(323F-390) hybrid under the control of the lacUV5 promoter in pPSV37 | This study |
| B24 | pP37-acrHpcrH-APB 369-popD | Plasmid encoding acrH, pcrH, popD and a aopB(1-368M)-popB(369I-390) hybrid under the control of the lacUV5 promoter in pPSV37 | This study |
| | pPGEH-acrHpcrH-popBD | Plasmid encoding acrH, pcrH, popB and popD under control of the PcrG promoter | This study |
| | pPGEH-acrHpcrH-Bmix-popD | Plasmid encoding acrH, pcrH, popD and a popB(1-114L)-aopB(108G-143K)-popB(150K-311V)-aopB(312S-330A)-popB(331Q-390) hybrid under control of the PcrG promoter | This study |

**Table S3: Primers**

| Primer Name | Sequence | Description |
|---|---|---|
| acrV-5 | CTTGTTGATCTGAGGAATCACGATGAGCGTTATCCCTGACTAC AACAG | 5' and 3' primers to amplify acrV |
| acrV-3 | AGGGGTCGGCTGGTTCATGGATACCTTCAAATAGCGCCAAGA ATGTCG | |
| acrVflank-5-2 | CTGTTGTAGTCAGGGATAACGCTCATCGTGATTCCTCAGATCA ACAAG | 5' and 3' primers to amplify acrV and flanking sequences |
| acrVflank-3-1 | CGACATTCTTGGCGCTATTTGAAGGTATCCATGAACCAGCCGA CCCCT | |
| acrV5R | AAAAAgaattcAAACATCAGGAGAAGGCAACCATCATGAGCGTTA TCCCTGACTACAAC | 5' and 3' primers to amplify acrV and add restriction sites EcoRI and XbaI |
| acrV3X | AAAAAtctagaTTAAATAGCGCCAAGAATGTCGCGCAT | |
| ANPCV-A5-2 | CAGCGCCTTGCGCTTGCCGTCCTGAAACGCCATCACCCCGGT AAAGAC | 5' and 3' primers to amplify N-terminal domain of acrV for fusion with pcrV to create Acollar |
| ANPCV-A3-1 | GTTGATCTGAGGAATCACGATGAGCGTTATCCCTGACTACAAC | |
| ANPCV-P5-2 | GTTGTAGTCAGGGATAACGCTCATCGTGATTCCTCAGATCAAC | 5' and 3' primers to amplify pcrV for fusion with N-terminal domain of acrV to create Acollar, which has AcrV(1-F143) fused to PcrV(Q124-end) |
| ANPCV-P3-1 | GTCTTTACCGGGGTGATGGCGTTTCAGGACGGCAAGCGCAAG GCGCTG | |
| pcrVAtip-5-2 | GTTGAAGGTCTGGCTGGATGCGGCCGACAGCGCGGCGTTGAT | pcrV primer for 5' flank fused to acrV tip (PcrV 1-158, AcrV 179-317, PcrV 251-294) |
| pcrVAtip-3-1 | GTGAGTGACCGCTCCCGTCCGCTGAACGACAAGGTCAACGAG | pcrV primer for 3' flank fused to acrV tip (PcrV 1-158, AcrV 179-317, PcrV 251-294) |
| acrV-tip5 | ATCAACGCCGCGCTGTCGGCCGCATCCAGCCAGACCTTCAAC | 5' and 3' primers for pcrV(acrVtip) fusion |
| acrV-tip3 | CTCGTTGACCTTGTCGTTCAGCGGACGGGAGCGGTCACTCAC | |
| pcrH-3SBS | TATATGTCGACCTCAGGATCCCTCAACTAGTTCAAGCGTTATC GGATTCGTATGCTCGATCCTTTC | pcrH 3' primer to introduce the speI site for new acrHpcrH constructs |
| acrHpcrH-5-2 | TGTCGGAAGGGGTCGGCTGGTTCATTGCTGATCGGGTTCGTC | 5' and 3' primers to concatenate acrH and pcrH, translationally coupled, and changes aopB start codon ATG->ACG |
| acrHpcrH-3-1 | GACGAACCCGATCAGCAATGAACCAGCCGACCCCTTCCGACA | |
| popB-5Spe | AAAAAactagtGTTTAAGGAGGAATAACCATGAATCCGATAACGC TTGAA | 5' and 3' primers to amplify popB with SpeI, XbaI, or BamHI restriction sites |
| popB-5Xba | AAAAAtctagaGTTTAAGGAGGAATAACCATGAATCCGATAACGC TTGAA | |
| popB3Bam | AAAAAggatccTCAGATCGCTGCCGGTCGGCTGGA | |
| aopB-5Xba | AAAAAtctagaGTTTAAGGAGGAATAACCATGAACCCGATCAGCA ATGAAAG | 5' and 3' primers to amplify aopB with XbaI or BamHI restriction sites |
| aopB-3Bam | TATATGGATCCTAAATAGCGGCGGGTCTGCGGGC | |
| popD5Bam | TATATggatccGTTTAAGGAGGAATAACCATGATCGACACGCAAT ATTCCCT | 5' and 3' primers to amplify popD with BamHI or SalI restriction sites |
| popD3Sal | AAAAAgtcgacCGCGCGGAGACGGCTCAGACCACT | |
| aopD-5Bam | TATATggatccGTTTAAGGAGGAATAACCATGATTAACAGTGATTA TGC | 5' and 3' primers to amplify aopD with BamHI or SalI restriction sites |
| aopD-3Sal | ATATAGTCGACTATGCAACACCGAATGCCGC | |
| APB115-3-1 | GCCGCGTTCGAGATTGAGTTGGCGAAAATCACCGGCGAAGTC GAG | 5' and 3' primers to fuse aopB 1-107(L) to popB 115(A)-390 |
| APB115-5-2 | CTCGACTTCGCCGGTGATTTTCGCCAACTCAATCTCGAACGCG GC | |
| APB169-3-1 | AAGAAAGCCGGTATTTTTGGCAAAATCTTTGGTTGGATCAGT | 5' and 3' primers to fuse aopB 1-161(G) to popB 169(K)-390 |
| APB169-5-2 | ACTGATCCAACCAAAGATTTTGCCAAAAATACCGGCTTTCTT | |
| APB183-3-1 | GCCATCGCCTCGGTCGTTATCGGCGCAATCATGGTGGCAACC | 5' and 3' primers to fuse aopB 1-175(I) to popB 183(G)-390 |
| APB183-5-2 | GGTTGCCACCATGATTGCGCCGATAACGACCGAGGCGATGGC | |
| APB251-3-1 | TCGCTGGTCGCGAGTTTTGGTGGCTCAGCGGTCGGCGGGCT G | 5' and 3' primers to fuse aopB 1-243(G) to popB 251(G)-390 |
| APB251-5-2 | CAGCCCGCCGACCGCTGAGCCACCAAAACTCGCGACCAGCG A | |
| APB298-3-1 | GTGACCGGGGCCGCTGCCAAGGTACTGAAACTCGGCGTGCA GGTTTCC | 5' and 3' primers to fuse aopB1-297(V) to popB 298(L)-390 |
| APB298-5-2 | GGAAACCTGCACGCCGAGTTTCAGTACCTTGGCAGCGGCCCC GGTCAC | |
| APB341 3-1 | cgtcattgaatgcgcaggctGACGTGCAGGAATCGCGTGC | 5' and 3' primers to fuse aopB1-332(A) to popB 333(D)-390 |
| APB341 5-2 | GCACGCGATTCCTGCACGTCagcctgcgcattcaatgacg | |
| APB344-3-1 | GAACAAGAGCGAGATGACCCAGCTGCAGGGCGTGATCGAGC GCCTG | 5' and 3' primers to fuse 1-343(Q) of aopB to 344(L)-390 of popB |
| APB344-5-2 | CAGGCGCTCGATCACGCCCTGCAGCTGGGTCATCTCGCTCTT GTTC | |

| Table S3: Primers | | |
|---|---|---|
| **Primer Name** | **Sequence** | **Description** |
| APB369-3 | CAGATCGCTGCCGGTCGGCTGGACAGGTTGTGCAGGGTTTCACCCTTGGCCTGGAGCATGGCGAAGATCATCTCCATGGTCTGCTGGAAC | 3' primer to fused 1-368(M) of aopB to 369(I)-390 of popB. Needs to be amplified with popB3Bam to include restriction site. |
| PAB40-3-1 | GTCTGCGGGTGGAGCGCCTGACGGCGCAGGCCAGTGGGCCAG | 5' and 3' primers to fuse fuse codon 1-40(L) of popB to codon 38(T)-390 of aopB |
| PAB40-5-2 | CTGGCCCACTGGCCTGCGCCGTCAGGCGCTCCACCCGCAGAC | |
| PAB114-3-1 | CAGGCGTTCGAGATCGAGCTGGGCAGGCTGACCAGCGATCTG | 5' and 3' primers to fuse popB 1-114(L) to aopB 108(G)-390 |
| PAB114-5-2 | CAGATCGCTGGTCAGCCTGCCCAGCTCGATCTCGAACGCCTG | |
| PAB164-3-1 | GCGAAAGAAGCCCAGAAGTCCGGTATTTTTGGCAAGATTTTTG | 5' and 3' primers to fuse popB1-164(S) to aopB 158(G)-390 |
| PAB164-5-2 | CAAAAATCTTGCCAAAAATACCGGACTTCTGGGCTTCTTTCGC | |
| PAB168-3-1 | CAGAAGTCCGGTCTGGCAGCCAAGATTTTTGGTTGGCTCAGT | 5' and 3' primers to fuse popB 1-168(A) to aopB 162(K)-390 |
| PAB168-5-2 | ACTGAGCCAACCAAAAATCTTGGCTGCCAGACCGGACTTCTG | |
| PAB250-5-2 | GCTAATGAGTGTGCCAATAGAGGAGCCGAAGCTGACCACCGCCGCCAG | 5' and 3' primers to fuse popB 1-250(G) to aopB 244(S)-390 |
| PAB250-3-1 | CTGGCGGCGGTGGTCAGCTTCGGCTCCTCTATTGGCACACTCATTAGC | |
| PAB297 3-1 | TCGCTGTCGCACgccaaagtcgcggagctggcagcccagttcacg | 5' and 3' primers to fuse popB 1-296(H) to aopB 270(A)-390 |
| PAB297 5-2 | gaactgggctgccagctccgcgactttggcGTGCGACAGCGATTGGC | |
| PAB330-3-1 | CAGGCGAAGGCCGCGAACCGCCAGGCTGATCTCGCGTTGAAC | 5' and 3' primers to fuse 1-330(R) of popB to 331(Q)-390 of aopB |
| PAB330-5-2 | GTTCAACGCGAGATCAGCCTGGCGGTTCGCGGCCTTCGCCTG | |
| PAPB 131-149 3-1 | cgcgcccgcgacctgaacgagaagaagatcggcgagaaccaggagAAGATCAGGGAGTCG | Forward primer to fuse aopB 142(E) to popB 150(K) |
| PAPB 131-149 5-2 | tcgttcaggtcgcgggcgcgcttgatgtcctccagCTTGAGTTTCTTCTG | Reverse primer to fuse popB 129(K) to aopB 123(L) |
| PAPB 312-330 3-1 | accaagaccgtgcagagcgtgaaggagagcgccagcctgaacgccCAGGCCGACGTGCAG | Forward primer to fuse aopB 330(A) to popB 331(Q) |
| PAPB 312-330 5-2 | ttcacgctctgcacggtcttggtcacgccgccgctGACGTCCAGGGTCAG | Reverse primer to fuse popB 311(V) to aopB 312(S) |
| APB 150 5-2 | GCTTCTTCCGACTCCCTGATCTTCTCCTGGTTCTCG | 5' and 3' primers to fuse aopB1-143(K) to popB 155(K) |
| APB 150 3-1 | CGAGAACCAGGAGAAGATCAGGGAGTCGGAAGAAGC | |
| PAB 131 5-2 | GCACGCTTGATATCCTCCAGCTTGAGTTTCTTCTGC | 5' and 3' primers to fuse popB1-131(T) to aopB 126(I) |
| PAB 131 3-1 | GCAGAAGAAACTCAAGCTGGAGGATATCAAGCGTGC | |
| PAB 151 5-2 | GCACTTTCTTCCGCCTCTTTGATCTTCTGCTGGTTGTCCTC | 5' and 3' primers to fuse popB1-151(I) to aopB 145K |
| PAB 151 3-1 | GAGGACAACCAGCAGAAGATCAAAGAGGCGGAAGAAAGTGC | |
| APB 331 3-1 | gtgaccaaaacggtccagtctgtgTTCCAGGCGAAGGCCGCGAACCGCCAG | 5' and 3' primers to fuse aopB1-322(V) to popB 323(F) |
| APB 331 5-2 | CTGGCGGTTCGCGGCCTTCGCCTGGAAcacagactggaccgttttggtcac | |
| dblatoOri | GACCCAAGTACCGCCACCTAAgctgaggCTGTCAGACCAAGTTTACTC | delete bla fragment and insert BbvCI site |
| pGtoRepA | CGTTGCCGGAGCCTGTCAGGCACGGTCGGGTTTTTGTCGTTTTCGCAGGCTTCGCTGCGTCACGTAGTGGTGCGTTTTTTG | replace lacUV5 promoter with pcrG promoter |
| dblatoGent | GAGTAAACTTGGTCTGACAGcctcagcTTAGGTGGCGGTACTTGGGTC | Primer for Gibson assembly of pPG-popD plasmid |
| dlacItoGent | CGGGTCTTGAGGGGTTTTTTGAGCGCAACGCAATTAATGTGAG | remove lacI gene |
| dlacItoTerm | CTCACATTAATTGCGTTGCGCTCAAAAAACCCCTCAAGACCCG | Primer for Gibson assembly of pPG-popD plasmid |
| pGtoPopD | CGACAAAAACCCGACCGTGCCTGACAGGCTCCGGCAACGTCGATCCCTACCATGGGCGACGGAATTCTTAGGAGGCGCC | Primer for Gibson assembly of pPG-popD plasmid |
| pGpolyFor | GGGGATCCTCTAGAGTCGACCTGCAGGCATGCAAGCTTTAGCATAACCCCTTGGGGCCTC | amplify pG-popD plasmid to make pG vector with polylinker from pPSV37 |
| pGpolyRev | CTCTAGAGGATCCCCGGGTACCGAGCTCCATATGAATTCCGTCGCCCATGGTAGGGATCG | |
